# How yeast cells find their mates

**DOI:** 10.1101/422790

**Authors:** Nicholas T. Henderson, Manuella R. Clark-Cotton, Trevin R. Zyla, Daniel J. Lew

## Abstract

Accurate detection of extracellular chemical gradients is essential for many cellular behaviors. Gradient sensing is challenging for small cells, which experience little difference in ligand concentrations on the up-gradient and down-gradient sides of the cell. Nevertheless, the tiny cells of the yeast *Saccharomyces cerevisiae* reliably decode gradients of extracellular pheromones to find their mates. By imaging the behavior of polarity factors and pheromone receptors during mating encounters, we found that gradient decoding involves two steps. First, cells bias orientation of initial polarity up-gradient, even though they have unevenly distributed receptors. To achieve this, they measure the local fraction of occupied receptors, rather than absolute number. However, this process is error-prone, and subsequent exploratory behavior of the polarity factors corrects initial errors via communication between mating partners. The mobile polarity sites convert the difficult problem of spatial gradient decoding into the easier one of sensing temporal changes in local pheromone levels.

## Introduction

Chemical gradients provide cells with critical information about their surroundings, allowing them to navigate via chemotropism (gradient-directed growth) or chemotaxis (gradient-directed migration). For example, axons steer their growth up gradients of netrin to form new synapses, social amoebae crawl up gradients of cAMP to aggregate into fruiting bodies, sperm swim up gradients of chemoattractants to find eggs, and neutrophils migrate up gradients of peptides shed by bacteria or cytokines secreted by other cells of the immune system to eliminate pathogens from mammalian tissues (Alvarez, et al., 2014; Swaney, et al., 2010; von Philipsborn and Bastmeyer, 2007). In each case, cells sense external signals via G-protein coupled receptors (GPCRs), leading to cytoskeletal reorganization that produces directional growth or movement (Insall, 2013).

The sequence of molecular events that transduce extracellular chemical signals to produce gradient-directed outputs is perhaps best understood in the genetically tractable budding yeast *Saccharomyces cerevisiae.* Yeast are non-motile unicellular fungi, and haploid yeast cells of mating type **a** can mate with haploids of mating type α to yield diploids. The haploids secrete peptide pheromones that bind GPCRs on cells of the opposite mating type (α-factor is sensed by Ste2 in **a** cells, and **a**-factor is sensed by Ste3 in α cells) (Wang and Dohlman, 2004). Pheromone-bound receptors activate heterotrimeric G-proteins to generate GTP-Gα and free Gβγ. Gβγ in turn recruits two key scaffold proteins, Ste5 and Far1, from the cell interior to the membrane (Nern and Arkowitz, 1999; Butty, et al., 1998; Pryciak and Huntress, 1998). Ste5 recruitment leads to activation of the MAPKs Fus3 and Kss1, which induce transcription of mating-related genes and arrest the cell cycle in G1 phase in preparation for mating (Pryciak and Huntress, 1998). Ste5-induced MAPK activation also promotes cytoskeletal polarization, but Far1 recruitment is required to orient the cytoskeleton towards the mating partner (Matheos, et al., 2004; Pryciak and Huntress, 1998; Valtz, et al., 1995). Far1 transduces the pheromone gradient by providing spatial information to the conserved Rho-family GTPase Cdc42, which is the master regulator of cell polarity in yeast (Bi and Park, 2012; Nern and Arkowitz, 1999; Butty, et al., 1998; Nern and Arkowitz, 1998).

To establish a polarized axis, Cdc42 becomes concentrated and activated at a site on the cell cortex referred to as a “polarity patch” (Bi and Park, 2012). The localized active Cdc42 then acts through formins to orient linear actin cables towards the site, and the cables deliver secretory vesicles that mediate local growth and fusion with a mating partner (Chen, et al., 2012; Liu, et al., 2012; Pruyne and Bretscher, 2000; Evangelista, et al., 1997). Polarity establishment is thought to involve a positive feedback loop whereby local GTP-Cdc42 promotes activation of further Cdc42 in its vicinity (Johnson, et al., 2011). Cdc42 is activated by the guanine nucleotide exchange factor (GEF) Cdc24 (Zheng, et al., 1994), which is recruited to the polarity patch by the scaffold protein Bem1, which is itself recruited to the patch by Cdc42 effectors, providing a mechanism for positive feedback (Kozubowski, et al., 2008). Cdc24 also binds directly to Far1, and the Gβγ-Far1-Cdc24 complex is thought to enhance GEF-mediated Cdc42 activation at sites with elevated levels of free Gβγ (Nern and Arkowitz, 1999; Butty, et al., 1998; Nern and Arkowitz, 1998). Mutations that disrupt Far1-Cdc24 binding do not affect polarity establishment *per se*, but they completely abolish the ability to properly orient polarity with respect to the pheromone gradient (Nern and Arkowitz, 1999; Butty, et al., 1998). Thus, Far1 provides a direct spatial connection between upstream receptor-pheromone binding and downstream Cdc42 activation, allowing the cells to exploit the pheromone gradient to find their partners.

Like other eukaryotic cells, yeast are thought to compare the ligand concentrations across the cell to determine the orientation of the gradient (Arkowitz, 2009). If the distribution of pheromone-activated receptors reflects the pheromone gradient, then Gβγ-Far1-Cdc24 complexes will be enriched up-gradient, spatially biasing activation of Cdc42 to kick off positive feedback at the right location for mating. However, the accuracy of such global spatial gradient sensing is limited by the small yeast cell size (∼4 μm diameter) (Berg and Purcell, 1977), and simulations constrained by experimental data on binding and diffusion parameters suggested that the process would be inaccurate (Lakhani and Elston, 2017). Indeed, when yeast are exposed to artificial, calibrated pheromone gradients, polarized growth often starts in the wrong direction (Moore, et al., 2008; Segall, 1993). Such cells can nevertheless correct initial errors by moving the polarity site (Dyer, et al., 2013).

Moving a Cdc42 patch that is constantly being reinforced by positive feedback seems counterintuitive, but time-lapse imaging revealed that the patch “wandered” around the cortex of pheromone-treated cells on a several-minute timescale (Dyer, et al., 2013). Wandering was dependent actin cables and vesicle traffic, which serves to perturb the polarity patch (Savage et al 2012; Dyer et al. 2013; McClure et al. 2015). New pheromone receptors are delivered to the polarity site, and after binding pheromone the receptors are rapidly internalized and degraded (Hicke, et al., 1998; Hicke and Riezman, 1996; Schandel and Jenness, 1994; Jenness and Spatrick, 1986). As a result, pheromone receptors and their associated G proteins become concentrated in the vicinity of the polarity site, generating a sensitized region of membrane that can detect the local pheromone concentration (McClure, et al., 2015; Suchkov, et al., 2010; Ayscough and Drubin, 1998). As the polarity site wanders around the cortex, this receptive “nose” would sample pheromone levels at different locations. Studies of cells treated with uniform pheromone concentrations showed that when pheromone levels are high, the patch stops moving (McClure, et al., 2015; Dyer, et al., 2013). In principle, this “exploratory polarization” mechanism can explain error-correction by positing that movement of the patch continues until cells sense high pheromone levels indicating that the patch is directed towards a mating partner (Hegemann and Peter, 2017).

The extent to which yeast cells rely on global spatial sensing to orient the formation of a polarity patch, versus exploratory polarization after the patch has formed, remains unclear. A recent study found that when cells were placed in an artificial pheromone gradient in a microfluidics device, initial patch formation was random with respect to the gradient, and orientation occurred by exploratory polarization and “local sensing” (Hegemann, et al., 2015). However, it is unclear whether similar results might apply to different pheromone gradients, or to more physiological conditions in which gradients are generated by mating partners.

To better understand how yeast actually locate their mating partners, we imaged mating events in mixed populations of **a** and α cells. We found evidence for both global spatial sensing and error correction by exploratory polarization. Encounters between partners were characterized by (i) rapid and non-random initial clustering of polarity proteins biased towards the partner; (ii) an “indecisive phase” in which dynamic polarity sites relocalized in an apparent search process; and (iii) a “committed phase” in which cells polarized stably towards mating partners, culminating in fusion. Transition from indecisive to committed behavior was associated with a rise in MAPK activity. Initial polarization was surprisingly accurate given that it occurred despite a highly non-uniform (and thus potentially misleading) distribution of receptors. We found that the variation in receptor density was corrected for via “ratiometric” sensing of the ratio of occupied vs unoccupied pheromone receptors across the cell (Bush, et al., 2016). In aggregate, our findings reveal how yeast cells can overcome the challenges imposed by small cell size and lack of cell mobility to locate mating partners.

## Results

### Indecisive and committed phases of mating cell polarization

To observe how cells find their mates, we mixed **a** and α cells expressing differently colored polarity probes, Bem1-GFP (α) and Bem1-tdTomato (**a**), and imaged them at 2 min resolution (Video 1). Fusion events were identified from movies and the cells were tracked back to their time of “birth” (the cytokinesis that preceded the mating event). Fig. 1A (top) illustrates selected frames from a representative mating cell. This cell formed a faint initial cluster of Bem1 just 4 min after birth (blue panel), which then fluctuated in intensity and moved erratically around the cell cortex for 34 min before stably polarizing adjacent to a mating partner (orange panel). After another 16 min the two cells fused, as seen by the mixing of red and green probes. We designate the time between initial cluster formation (T_ic_) and stable polarization (T_p_) as the “indecisive phase”, reflecting the erratic behavior of the polarity probe. We designate the time between stable polarization (T_p_) and fusion as the “committed phase” of mating, reflecting the strong and stably located polarity site.

**Figure 1.**
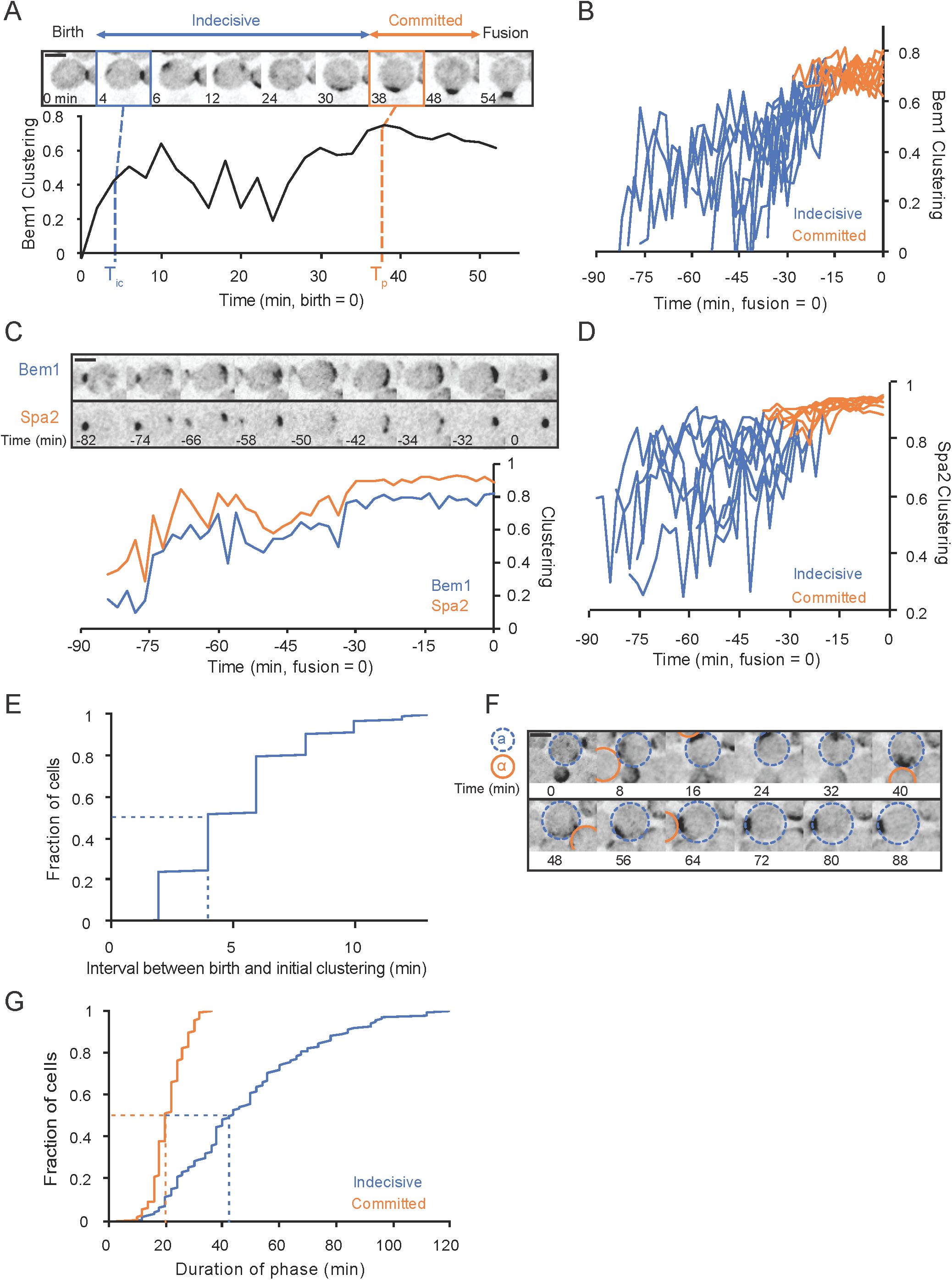
Mating yeast display distinct indecisive and committed stages of polarity behavior. (**A**) Localization of Bem1-GFP in a representative mating cell. Top: Inverted maximum z-projection images of Bem1-GFP at selected time points, with birth (cytokinesis) designated as 0 min. A weak Bem1 cluster appears 4 min (blue box, T_ic_ – time of initial clustering). The cluster moves and fluctuates in intensity during an “indecisive phase” until 38 min (orange box, T_p_ – time of polarization), when it strengthens and remains stationary during a “committed phase” until fusion occurs at 54 min. Bottom: quantification of Bem1 clustering in the same cell (see methods). (**B**) Bem1 clustering plotted for 10 representative mating cells as in (A). In this panel, the time of fusion was designated as 0 min and the timeline extends back to the time of cell birth. Color switches from blue to orange at Tp. (**C**) Localization of Bem1-GFP and Spa2-mCherry in a mating cell from birth (−82 min) to fusion (0 min). Top: Inverted maximum z-projection images of the indicated probes. Bottom: quantification of Bem1 and Spa2 clustering in the same cell. (**D**) Spa2 clustering in 10 representative cells, displayed as in (B). (**E**) The cumulative distribution (n=150) of the interval between birth and T_ic_ in mating cells. (**F**) Localization of Bem1-tdTomato in a “serial dating” cell. Inverted maximum z-projection images of selected time points displaying a MAT**a** cell transiently orienting polarity towards 4 different MATα cells before committing to the 1st. The MAT**a** cell is circled in blue, and each sequential partner is circled in orange. (**G**) The cumulative distribution (n=150) of the duration of the indecisive phase (blue) and the committed phase (orange). Dashed lines indicate median. Scale bar, 3 μm. Strains: DLY12943, DLY7593 (A, B, E-G), DLY21379 (C, D).

To quantify the degree of Bem1 polarization, we used a metric that uses the pixel intensity distribution within the cell to assess the degree of signal clustering (hereafter, “clustering”: see methods). Fig. 1A (graph) illustrates that Bem1 clustering fluctuated during the indecisive phase but remained high during the committed phase. This pattern was characteristic of mating cells (Fig. 1B), supporting the idea that mating involves a two-stage process. A similar two-stage process was observed for cells expressing fluorescent versions of Spa2, a polarisome component that binds and helps to localize the formin Bni1 (Video 2) (Pruyne, et al., 2004; Fujiwara, et al., 1998; Sheu, et al., 1998). Analysis of cells expressing both Bem1 and Spa2 probes revealed that although (as described previously) Spa2 clusters were more tightly focused than Bem1 clusters, the probes clustered, dispersed, and moved together (Fig. 1C). As for Bem1, Spa2 clustering fluctuated during the indecisive phase and remained stably high during the committed phase (Fig. 1D). We conclude that cells undergo a reproducible pattern of polarization during mating, with sequential indecisive and committed phases.

The earliest observable clustering of polarity factors occurred shortly after birth (Fig. 1E: median time 4 min after initiating cytokinesis). This initial clustering was usually weak and frequently at a different location than that of the final stable polarization (see below). During the ensuing indecisive phase, cells appeared to search for mating partners, often assembling polarity clusters adjacent to different potential partners before settling at a final location (Fig. 1F). The duration of the indecisive phase (Fig. 1G: median 42 min) was very variable, ranging from 10 to 120 min. This is consistent with a search process that would take a variable amount of time depending on the availability and proximity of potential mating partners. In contrast, the subsequent committed phase was consistently about 20 min (Fig. 1G), which we speculate is the time required to remodel the local cell walls to allow for cell fusion.

### Commitment is synchronous for both partners

In our protocol, cells of each mating type are proliferating asynchronously before they are abruptly mixed. Thus, in a large majority of cases, one cell of each mating pair is born (i.e. enters G1 phase) before the other. Nevertheless, fusion is a unitary event that occurs at the same time for both. This means that the “first-born” partner must extend one or both phases of polarization while the “second-born” partner completes the previous cell cycle and “catches up” (Fig. 2A). Does the first-born locate and commit to its partner first, and then wait (Fig. 2A, top), or does the first-born remain indecisive until the second-born has caught up (Fig. 2A, bottom)? We found no difference in the average duration of the committed phase between first and second-born cells (Fig. 2B), and partners in each individual mating pair generally committed at nearly the same time (Fig. 2C). Conversely, the indecisive phase was significantly longer in first-born cells (Fig. 2D), suggesting that first-born cells remain indecisive while second-born cells complete the cell cycle, and that cells only polarize stably towards partners that are in G1 (Fig. 2A, bottom).

**Figure 2.**
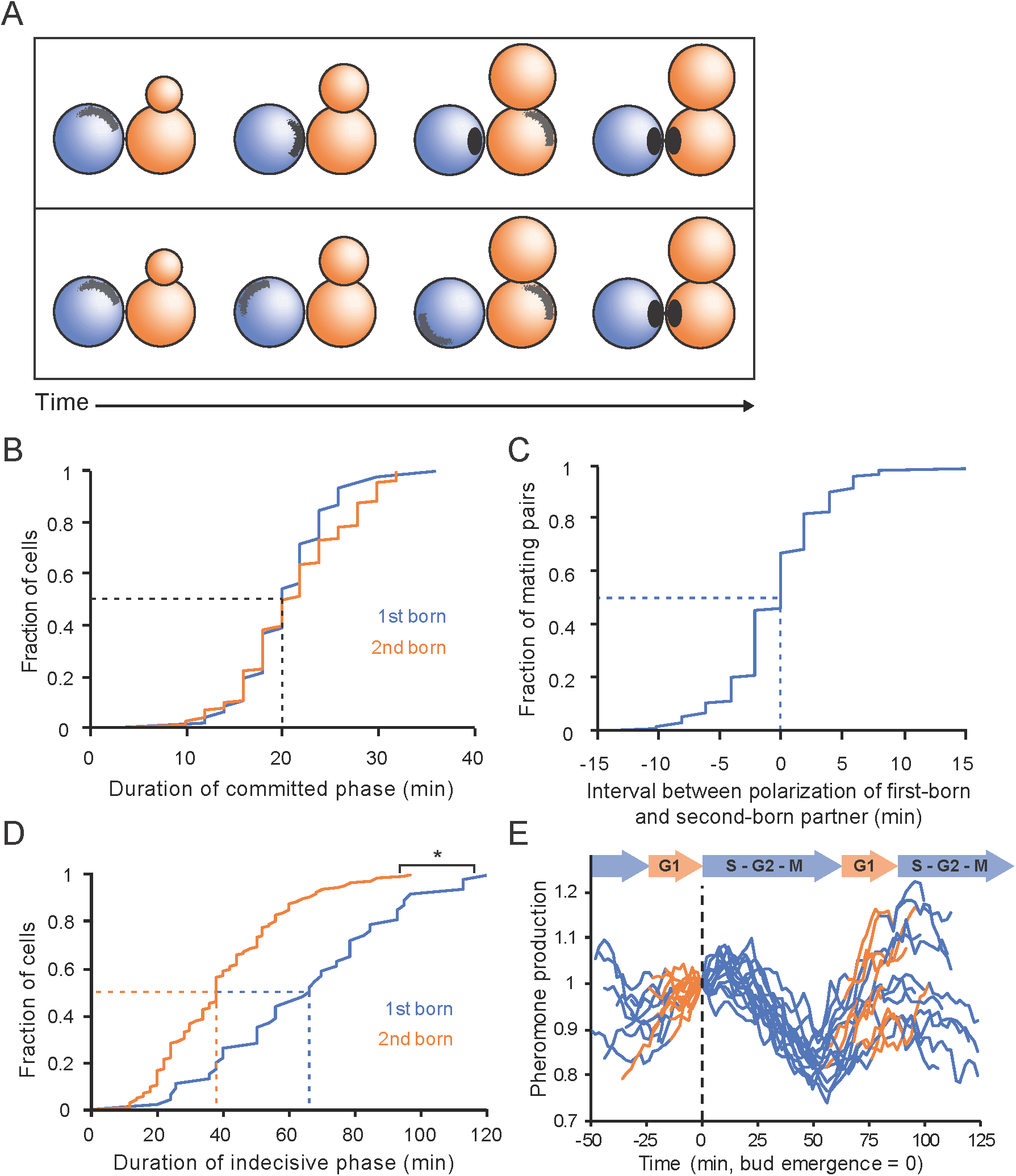
Synchronous commitment by both partners. (**A**) Two hypotheses for polarization timing in mating partners that “meet” when they are in different stages of the cell cycle. Top: The first-born cell (blue) locates the partner (orange) while it is still completing the cell cycle. The first-born cell polarizes and waits during an extended committed phase for its partner to catch up. Bottom: The first-born cell cannot locate its partner until the partner enters G1 phase. It remains in an extended indecisive phase until the partner enters G1, after which both cells locate one another and polarize simultaneously. (**B**) Cumulative distribution of the interval between stable polarization and fusion (i.e. duration of the committed phase) in first-born (blue, n=46) and second-born cells (orange, n=104). (**C**) Cumulative distribution of the interval between when the first-born cell polarizes and when the second-born cell polarizes (i.e. T_p2_ - T_p1_) (n=104). (**D**) Cumulative distribution of the interval between initial clustering and commitment (i.e. duration of the indecisive phase) in first-born (blue, n=46) and second-born cells (orange, n=104) (* two sample Kolmogorov-Smirnov [KS] test, p < 0.05). (**E**) Pheromone synthesis is high in G1 and decreases as cells enter the cell cycle. Cells harboring the reporter sfGFP under control of the MFα1 promoter were imaged for 150 min at 2 min resolution. Reporter fluorescence was normalized to the value at the time of first bud emergence. Bud emergence designated as 0 min. Curves were colored orange from birth to bud emergence (G1 phase), and blue from bud emergence to birth (S, G2, and M phase). Dashed lines indicate median (B-D) or times of bud emergence (E). Strains: DLY12943, DLY7593 (B-D), DLY22883 (E).

Synchronous commitment implies that there is some communication between partners that occurs only when both are in G1 phase of the cell cycle. As the only known mode of communication is via the secretion of pheromones, the simplest hypothesis to explain why commitment must wait until both cells are in G1 would be that pheromone secretion changes when cells enter G1. To assess the rate of pheromone synthesis, we introduced a fluorescent reporter whose production was driven by the major α-factor gene (MFα1) promoter (Achstetter, 1989; Singh, et al., 1983). Reporter signal fluctuated regularly through the cell cycle, rising in G1 and falling (due to dilution) after bud emergence (Fig. 2E). This result suggests that first-born cells would detect lower levels of pheromone until their partners entered G1, and that stable polarization towards a partner (commitment) may be triggered by increased pheromone signaling.

### Commitment coincides with an increase in MAPK activity

One consequence of pheromone signaling is the activation of the mating MAPKs Fus3 and Kss1 (Wang and Dohlman, 2004). To monitor MAPK activity in mating cells, we introduced a recently developed single-cell MAPK sensor (Durandau, et al., 2015) into our strains together with the Spa2 probe. The MAPK sensor is a fluorescent probe that moves from the nucleus to the cytoplasm when it is phosphorylated by active MAPK. In the absence of pheromone, the sensor was predominantly nuclear, although the nuclear to cytoplasmic ratio fluctuated somewhat through the cell cycle, peaking during anaphase (Fig. 3A and video 3). In a mating mix, the sensor distribution became uniform prior to fusion, reflecting an increase in MAPK activity (Fig. 3B and video 4). To quantify the degree of nuclear concentration of the MAPK sensor, we measured the coefficient of variation (CV) in pixel intensity across the cell. When the probe is nuclear, the bright nuclear and dim cytoplasmic pixels yield a high CV, but when the probe distribution is uniform there is a low CV. We found considerable cell-to-cell variability in this signal, which could be largely accounted for by differences in the level of expression of the probe (Fig. S1A, B). This variability could be reduced by normalizing the CV to the maximum and minimum CV for each cell, and we developed a MAPK activity metric based on the normalized CV of the probe (Fig. S1C).

**Figure 3.**
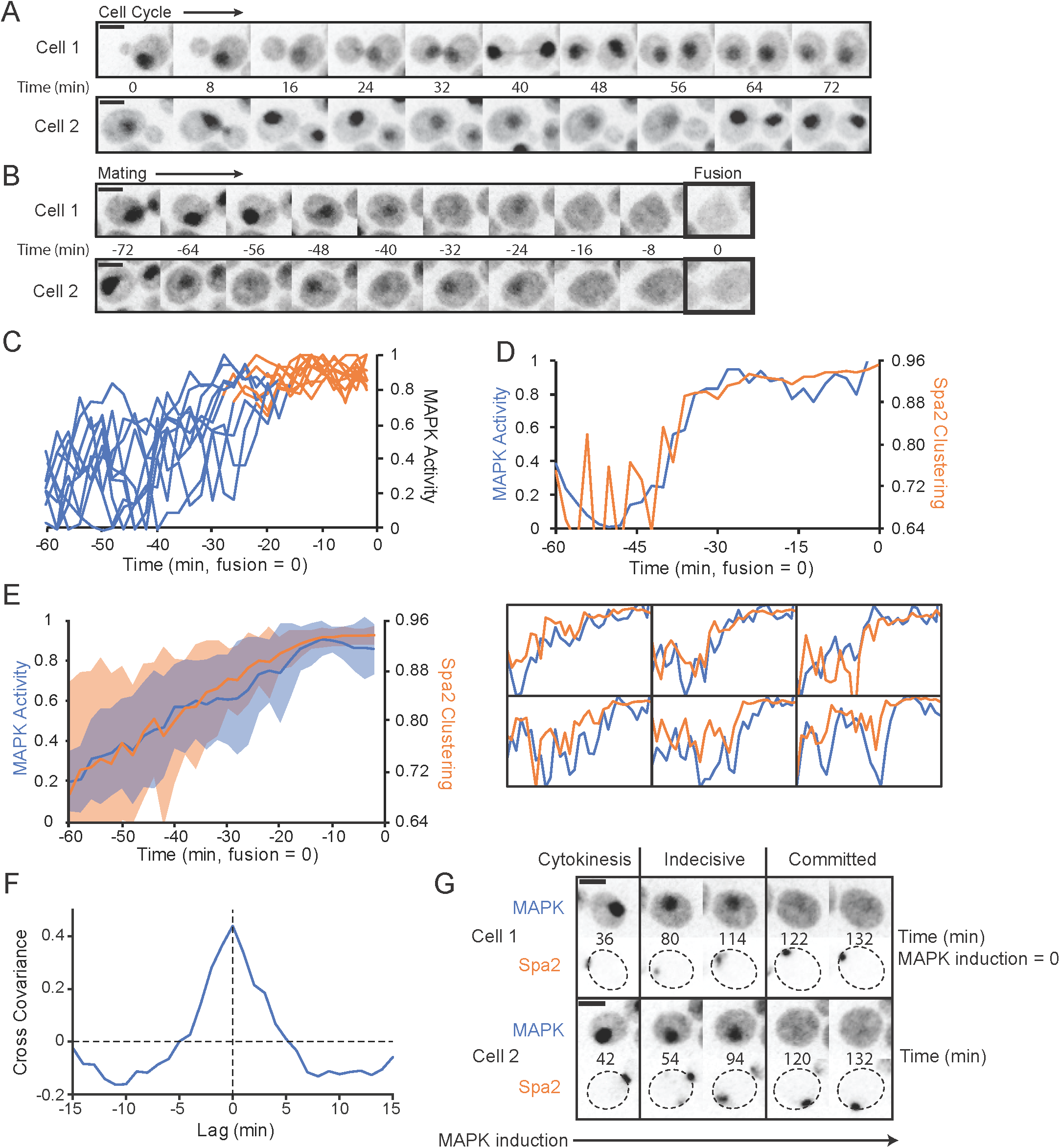
Commitment coincides with an increase in MAPK activity. (**A**) Localization of MAPK activity sensor varies through the cell cycle. Inverted maximum z-projection images of the sensor Ste7_1-33_-NLS-NLS-mCherry in two representative vegetatively growing cells. (**B**) The sensor is exported from the nucleus in response to MAPK activation. MAT**a** cells harboring Ste7_1-33_-NLS-NLS-mCherry were mixed with MATα cells and imaged as in (A). Two representative mating cells illustrated from birth to fusion (0 min = fusion). (**C**) MAPK activity calculated from sensor distribution (see methods) in the 60 min prior to fusion. The transition from the indecisive phase (blue) to the committed phase (orange) was determined from a Spa2-GFP probe in the same cells. (**D**) Top: MAPK activity (blue, as in C) and Spa2 clustering (orange, as in Fig. 1F) in a representative mating cell. Bottom: six other cells plotted as above. (**E**) Average MAPK activity (blue) and Spa2 clustering (orange) during the 60 min prior to fusion (n=23 cells). Shaded regions represent standard deviations. (**F**) Cross-covariance of MAPK activity and Spa2 clustering during the indecisive phase (window from 60 minutes before fusion to 30 minutes before fusion) in mating cells (n=23 cells). Lag represents the time by which the Spa2 clustering data was shifted forward in time relative to the MAPK activity data. 1 = perfect cross-covariance (i.e. auto-covariance = 1 when lag = 0) (**G**) Cells harboring P_GAL1_-Ste5-CTM allow MAPK induction by β-estradiol without pheromone treatment. The MAPK sensor Ste7_1-33_-NLS-NLS-mCherry and Spa2-GFP were imaged following β-estradiol treatment and inverted maximum z-projection images of selected time points showing Spa2 neck localization during cytokinesis, the indecisive behavior upon intermediate MAPK activation, and committed behavior following high MAPK activation in two representative cells. Scale bar, 3 μm. Strains: DLY22259 (A-F), DLY22764 (G).

In mating cells, MAPK activity fluctuated but then climbed to a plateau about 20 min prior to fusion (Fig. 3C). As this was similar to the clustering behavior of polarity probes, we directly compared MAPK activity with Spa2 clustering in individual mating cells (Fig. 3D). These measures aligned well with one another in most cells, with both Spa2 clustering and MAPK activity fluctuating during the indecisive phase before rising to a stable plateau during the committed phase (Fig. 3D, E). A cross-correlation analysis of Spa2 clustering and MAPK activity during the indecisive phase revealed that they fluctuated in tandem (Fig. 3F). This correlation suggested that MAPK activity might promote stable polarization, or that polarization might lead to an increase in MAPK activity, or both.

To more directly ask whether an increase in MAPK activity promotes stable polarization, we induced MAPK activity in the absence of pheromone by expressing a membrane-tethered version of the MAPK scaffold Ste5 (Pryciak and Huntress, 1998). As MAPK activity increased and the MAPK sensor exited the nucleus, Spa2 switched from faint and mobile clustering to become strongly polarized (Fig. 3G). The timing varied from cell to cell, but all cells with induced MAPK eventually arrested and formed strong polarity patches (data not shown). Thus, elevated MAPK signaling is sufficient to induce polarization.

### Relation between pheromone sensing and polarization

Our findings suggest that at some point during the indecisive phase, MAPK activity rises, triggering the strong and stable polarization characteristic of the committed phase. But what would make the MAPK activity increase at that point? One possibility is that pheromone secretion is tied to polarity site behavior, so that pheromone is secreted from transient and erratically changing locations during the indecisive phase. Prior work has shown that cells have a sensitized “nose” enriched in receptors and G proteins surrounding the polarity site (McClure, et al., 2015; Suchkov, et al., 2010), so if two cells happen to orient their polarity sites towards each other during the indecisive phase, one cell would emit pheromone in the immediate vicinity of receptors on the partner cell. That would produce a higher pheromone signal than when polarity clusters point away from each other (Fig. 4A). To visualize the location of α-factor secretion, we imaged Sec4-GFP, a Rab GTPase highly concentrated on the secretory vesicles that deliver α-factor to the cell surface (Mulholland, et al., 1997; Walch-Solimena, et al., 1997). Sec4 accumulated in regions enriched for Bem1, during the indecisive phase as well as the committed phase of polarization (Fig. 4B).

**Figure 4.**
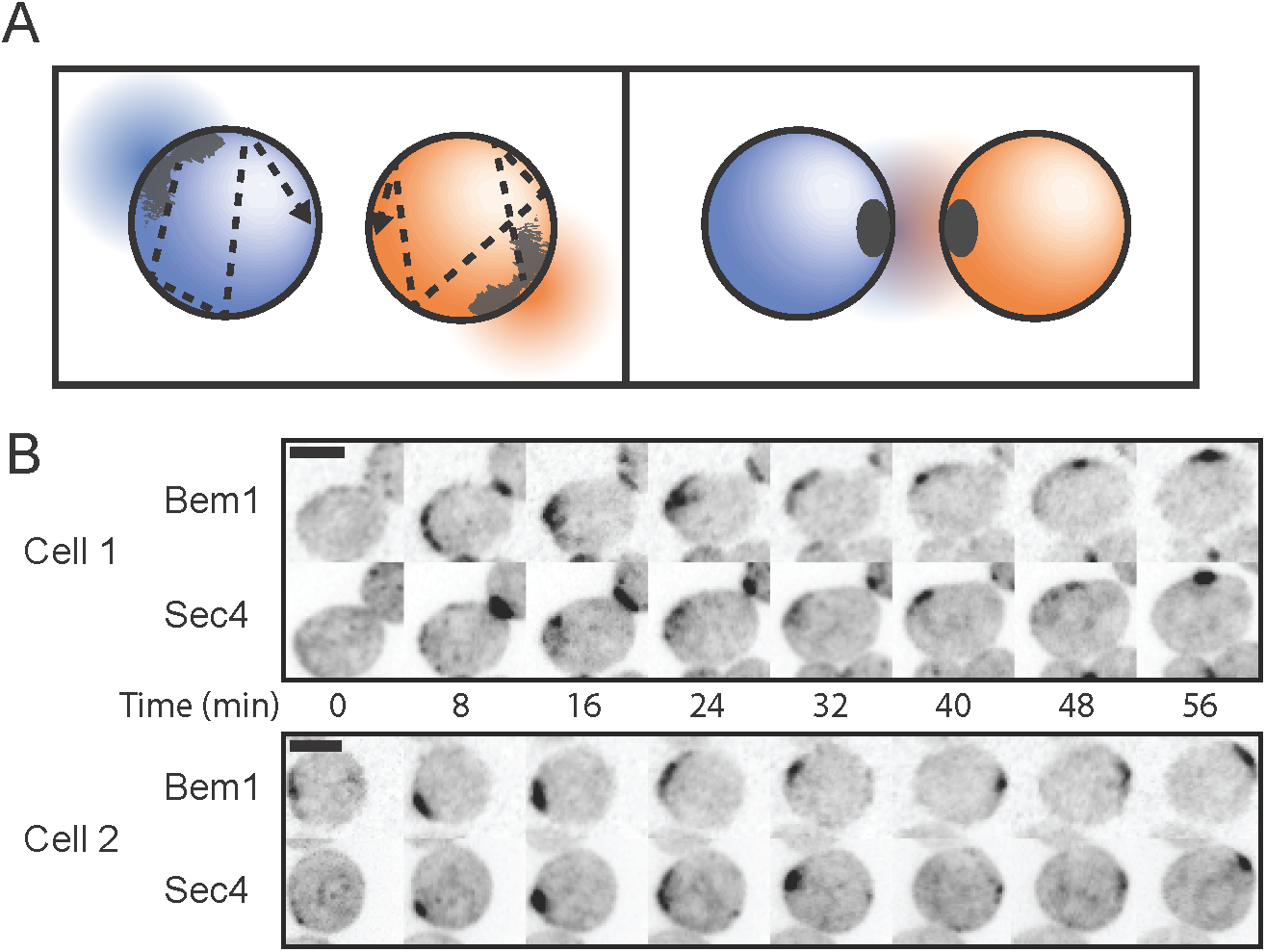
Secretory vesicles colocalize with the polarity cluster. (**A**) Hypothesized pheromone secretion during indecisive (left) and committed (right) phases. When clusters are adjacent, local pheromone concentration would be higher, triggering commitment. (**B**) MAT**a** cells harboring Bem1-tdTomato and the secretory vesicle marker GFP-Sec4 were mixed with MATα cells and imaged as in Fig. 1A. Inverted maximum z-projection images at 8 min intervals show colocalization of Bem1 and Sec4 during the indecisive phase. Scale bar, 3 μm. Strain: DLY13771.

Putting together our findings thus far, we propose that mating cells undergo the following sequence of events. As cells undergo cytokinesis, newborn cells in G1 phase detect enough pheromone in their surroundings to arrest the cell cycle and initiate a weak and mobile level of polarization. Cells in G1 also increase their rate of pheromone production, signaling to potential partners that they are ready to mate. As mobile polarity clusters explore the surrounding pheromone landscape during the indecisive phase, they also locally secrete pheromones to be sensed by their partners. When potential partners orient their polarity clusters towards each other, both cells perceive higher pheromone concentrations, leading to a simultaneous rise in MAPK activity in each partner. Higher MAPK activity leads to stronger polarity, and when MAPK activity crosses some threshold, the polarity clusters stop moving, now properly facing their partners. This leads to sustained high MAPK signaling during the committed phase, maintaining polarity until fusion can occur.

### Gradient sensing before initial polarity clustering

The view of the mating process outlined above proposes an important role for polarity clusters in tracking pheromone gradients to locate partners, as recently suggested by other studies which noted mobile polarity sites in cells exposed to uniform pheromone (McClure, et al., 2015; Dyer, et al., 2013) or artificial pheromone gradients (Hegemann, et al., 2015). This idea differs markedly from the traditional view, in which initially unpolarized G1 cells first sense the pheromone gradient and only then polarize, generally in the right direction (Ismael and Stone, 2017; Ismael, et al., 2016; Suchkov, et al., 2010; Arkowitz, 2009). These views are not mutually exclusive, and it could be that significant gradient sensing takes place prior to the initial clustering of polarity factors. Indeed, we found that in our mating mixtures, cells biased the locations of their initial clusters towards their eventual mating partners (Fig. 5A), suggesting that a form of global spatial gradient sensing occurs in the few minutes between cell birth and initial clustering. In contrast, we found no bias towards the previous cytokinesis site (neck) under our conditions (Fig. 5B). The directional bias towards partners was similar in first-born and second-born cells (Fig. 5C). Among second-born cells, those that formed their initial clusters within 60° of their partners took less time to commit than those whose initial clusters were less well-oriented (Fig. 5D: median indecisive phase duration 32 min vs 48 min). Thus, gradient sensing before polarity cluster formation can shorten the search for a partner.

**Figure 5.**
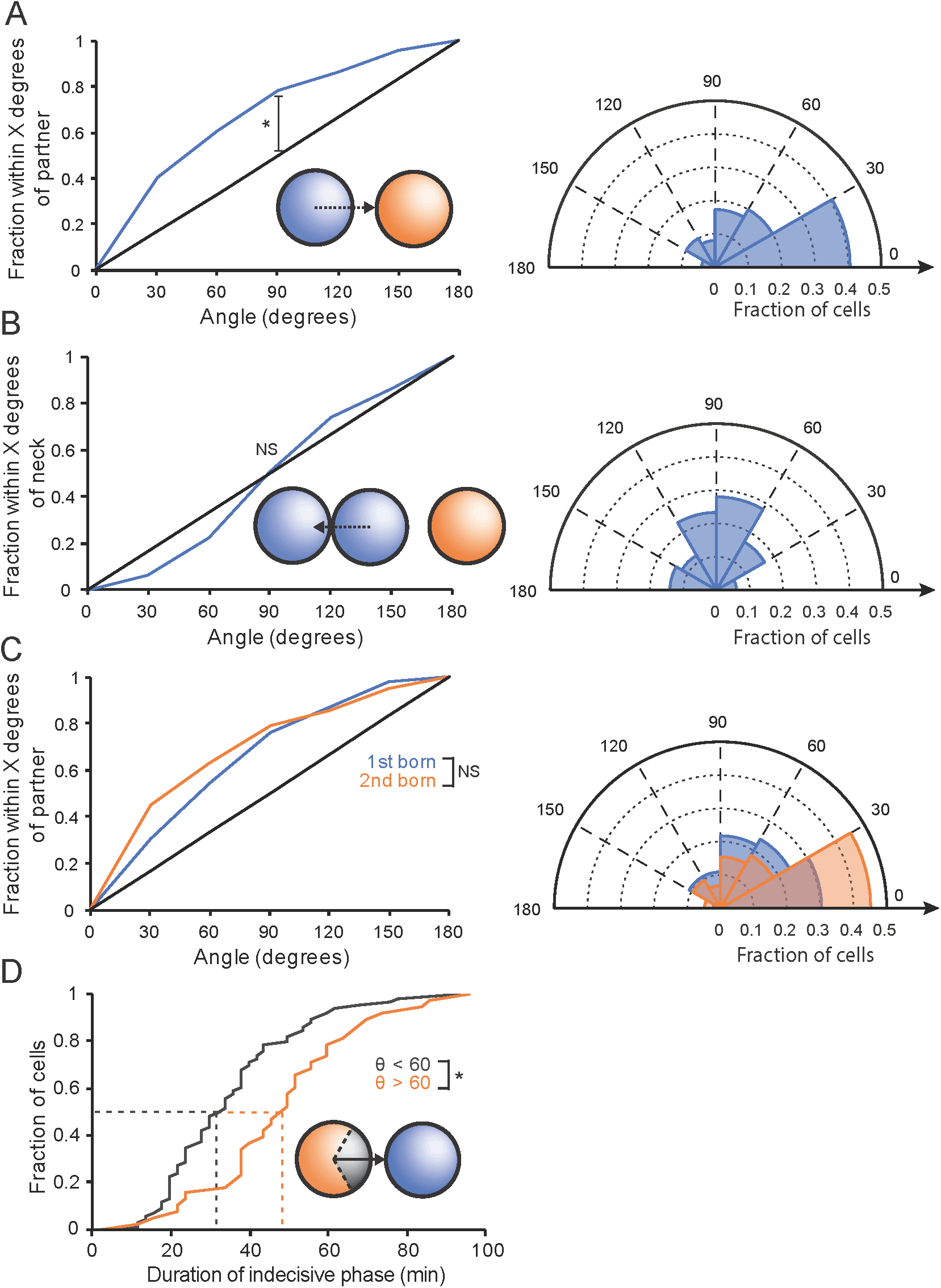
Non-random initial clustering of polarity factors. (**A**) Orientation with respect to partner. Left: cumulative distribution of initial Bem1 cluster location (angle) relative to the mating partner (n=150). 0 degrees = cluster formation directly adjacent to eventual mating partner, as shown in inset. Black line: hypothetical random distribution (* KS test, p < 0.05). Right: polar histogram displaying the same data. (**B**) Orientation with respect to neck. Left: cumulative distribution of initial cluster location relative to the site of cytokinesis (n=150, KS test, not significant). Right: polar histogram displaying the same data. (**C**) Left: cumulative distribution of initial cluster location relative to the mating partner, plotted separately for first-born (blue, n=46) and second born (orange, n=104) cells (two sample KS test, not significant). Right: polar histogram of the same data. (**D**) Cumulative distribution of the duration of the indecisive phase in second-born cells, plotted separately for cells in which the initial cluster formed within 60° of the mating partner (θ < 60°, orange, n=66), and cells in which the initial cluster formed greater than 60° from the partner (θ > 60°, blue, n=38)(* two sample KS test, p < 0.05). Inset: diagram displaying the two groups of cells. Strain: DLY12943, DLY7593.

### Non-uniform pheromone receptor distribution

How would spatial gradient sensing occur? Studies in other model systems like *Dictyostelium discoideum* indicated that receptors and their coupled G proteins were distributed uniformly around the cell surface, with active G proteins reflecting the external ligand gradient (Janetopoulos, et al., 2001; Jin, et al., 2000). Receptor distribution has been harder to assess in yeast cells, for technical reasons stemming from the rapid secretion and recycling of receptors (Suchkov, et al., 2010). Transit of pheromones and pheromone receptors through the secretory pathway is rapid (5-10 min)(Losev, et al., 2006; Govindan, et al., 1995). In the presence of α-factor, Ste2 is then endocytosed on a 10-min timescale and delivered to the vacuole for degradation (Hicke, et al., 1998; Hicke and Riezman, 1996; Schandel and Jenness, 1994; Jenness and Spatrick, 1986). As GFP maturation occurs on a 30-min timescale (Iizuka, et al., 2011; Gordon, et al., 2007), much of the GFP-tagged receptor at the cell surface is not yet fluorescent. Moreover, the GFP moiety survives intact in the vacuole following receptor degradation, yielding a high vacuolar fluorescence signal. To partially resolve these issues, we tagged Ste2 with sfGFP, which matures on a 6-min timescale (Khmelinskii, et al., 2012). Although bright vacuoles remained, the surface Ste2-sfGFP was clearly visible (Fig 6A), allowing us to assess Ste2 distribution. In cells that were not exposed to α-factor, Ste2 distribution varied throughout the cell cycle, accumulating in the bud (enriched at the tip) and depleted in the mother during bud growth, and then accumulating at the neck during cytokinesis (Fig. 6B). G1 cells displayed quite variable Ste2 distributions, ranging from nearly uniform to highly polarized (Fig. 6C: left). Quantification of surface Ste2 distribution revealed a 3-fold difference (on average) in Ste2 concentration from one side of the cell to the other (Fig. 6C: right).

**Figure 6.**
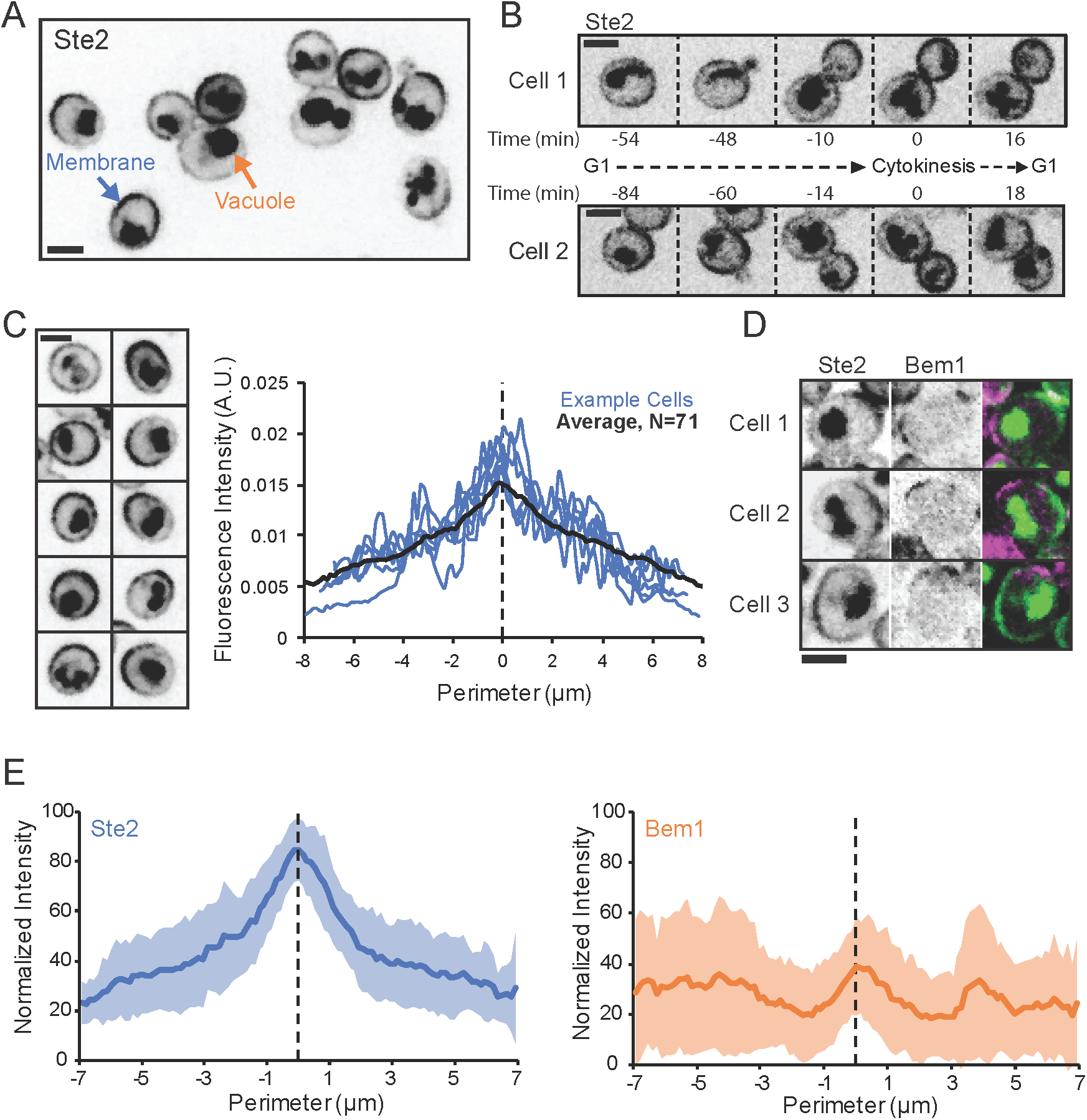
Pheromone receptor distribution prior to pheromone exposure. (**A**) Single-plane inverted image of vegetatively growing cells expressing Ste2-sfGFP. Membrane signal (blue arrow) is Ste2-sfGFP, but vacuole signal (orange arrow) is probably sfGFP cleaved from Ste2-sfGFP after internalization. (**B**) Time series of two representative cells displaying the Ste2 distribution through the cell cycle. (**C**) Left: Representative G1 cells displaying Ste2 distributions ranging from almost uniform (top left) to very asymmetric (bottom right). Right: quantification of Ste2-sfGFP membrane distribution in G1 cells. Individual linescans (examples in blue) were normalized to have the same total fluorescence and centered based on the maximum of a smoothed spline fit. Black line, average (n=71). (**D**) Initial Bem1 clusters sometimes form in areas depleted of receptor. Single-plane Ste2-sfGFP images (left), maximum projection Bem1-tdTomato images (middle), and both overlaid (right, Bem1 = magenta, Ste2 = green, both = white) from three example cells at the time of initial clustering. (**E**) Bem1 initial cluster formation is random with respect to Ste2 distribution. Left: averaged Ste2-sfGFP distribution (shaded region, standard deviation) at the time of initial clustering (n=33). Right: averaged Bem1-tdTomato distribution at the time of initial clustering, centered on the peak of the Ste2-sfGFP distribution (n=33). Bem1 linescans acquired from maximum projection images of the same cells. Scale bar, 3 μm. Strains: DLY20713 (A-C), DLY22243 (D, E).

The non-uniform receptor distribution poses a significant problem for accurate gradient sensing, because cells would be preferentially sensitive to pheromone on the side where receptors are enriched, which would not necessarily correspond to the side facing a mating partner. To directly observe the relationship between Bem1 clustering and Ste2 distribution, we imaged MAT**a** cells carrying both Ste2-sfGFP and Bem1-tdTomato, mixed with MATα cells in mating reactions. If receptor density impacts the location of initial clustering, we would expect that Bem1 clustering would occur preferentially on the side with higher Ste2 signal. Individual cells clustered Bem1 at various different locations relative to the Ste2 distribution, and in several cells the initial Bem1 cluster formed adjacent to a mating partner even though Ste2 was concentrated at the opposite end of the cell (Fig. 6D). Averaging revealed no clear spatial correlation between the location of Bem1 initial clustering and the Ste2 distribution (Fig. 6E). We conclude that cells are able to perform a surprisingly accurate degree of gradient sensing prior to polarization, despite having non-uniform receptor distributions.

### Effect of changing receptor distribution on gradient sensing

To probe the degree to which receptor distribution influences the accuracy of gradient sensing, we sought to manipulate receptor distribution. Ste2 distribution reflects a dynamic balance between polarized secretion of new Ste2, slow diffusion at the plasma membrane, and retrieval by endocytosis (Suchkov, et al., 2010; Valdez-Taubas and Pelham, 2003). Endocytosis is more rapid for ligand-bound Ste2 (which undergoes phosphorylation and ubiquitination) than for unbound Ste2 (which is endocytosed at a slower basal rate)(Hicke, et al., 1998; Terrell, et al., 1998; Hicke and Riezman, 1996). To manipulate Ste2 distribution, we used Ste2 mutants that either lacked endocytosis signals (Ste2^7XR-GPAAD^, allowing accumulation all over the membrane) (Ballon, et al., 2006; Terrell, et al., 1998) or had a constitutively active strong endocytosis signal (Ste2^NPF^ yielding a highly polarized distribution with a bias toward the mother-bud neck)(Tan, et al., 1996) (Fig. 7A, B). As endocytosis is needed for Ste2 degradation, Ste2^7XR-GPAAD^ was more abundant than Ste2 or Ste2^NPF^ (Fig. 7C), and in halo assays cells expressing Ste2^NPF^ were slightly less sensitive to pheromone while cells expressing Ste2^7XR-GPAAD^ were more sensitive to pheromone (Fig. 7D). Correspondingly, in mating mixes cells with Ste2^NPF^ sometimes re-entered the cell cycle despite being adjacent to potential partners, while cells with Ste2^7XR-GPAAD^ were more likely to remain arrested and mate (Fig. S2). This was reflected in the duration of the indecisive phase, which we quantified among all cells that were born and remained immediately adjacent to a G1 cell of opposite mating type until they either mated or budded (Fig. 7E). Cells that budded instead of committing to a partner were recorded as never entering the committed phase. Among the cells that successfully mated, indecisive phases had similar durations, suggesting that indecisive phase dynamics were unaffected by the changes in receptor distribution.

**Figure 7.**
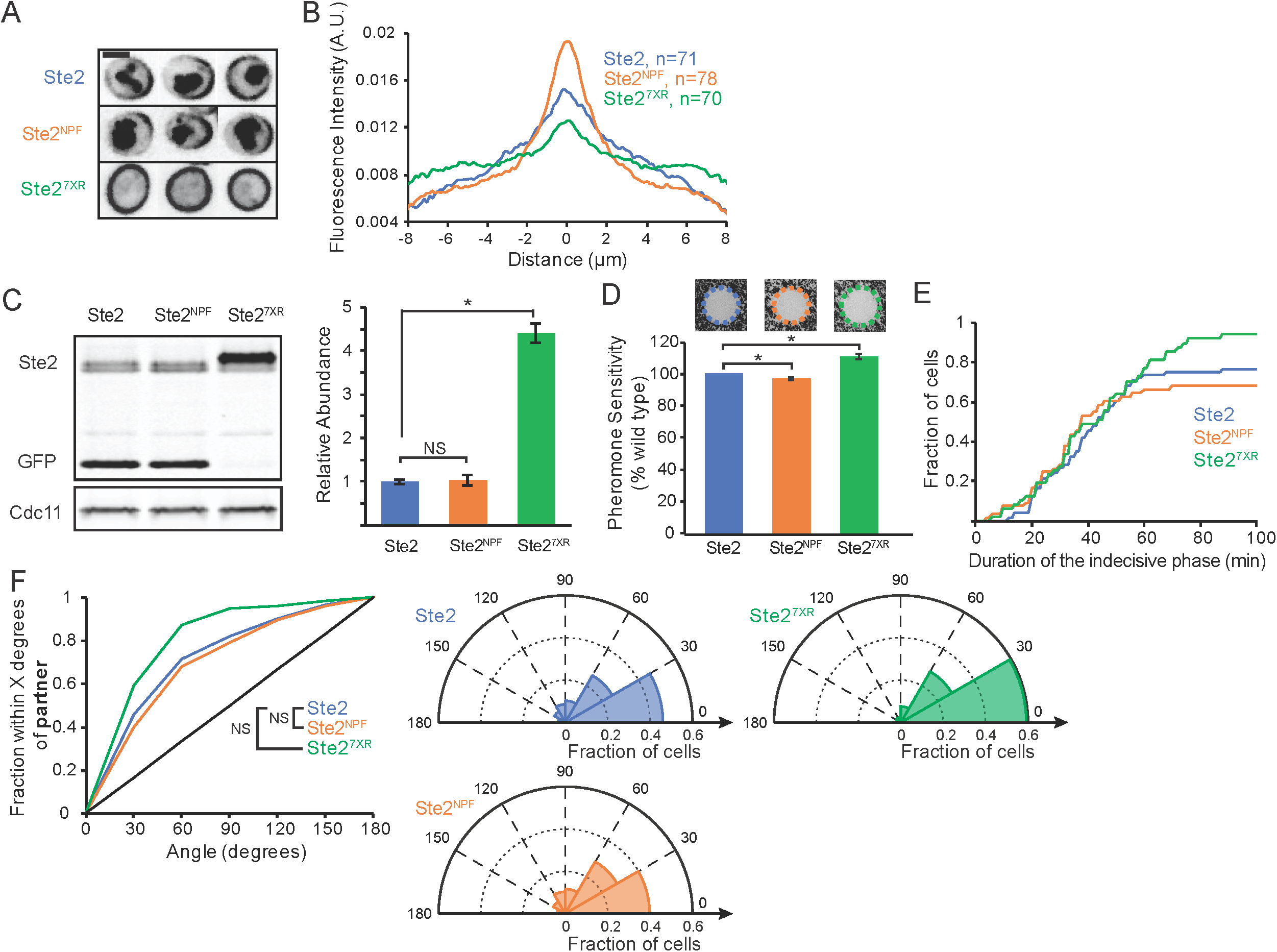
Effect of receptor distribution on the accuracy of initial clustering. (**A**) Single-plane inverted images of Ste2-sfGFP (top), Ste2^NPF^-sfGFP (middle), and Ste2^7XR-GPAAD^-sfGFP (bottom) in representative G1 cells. Ste2^7XR-GPAAD^-sfGFP displayed stronger fluorescence. As a result, the brightness and contrast of the Ste2^7XR-GPAAD^-sfGFP images have been scaled differently for clear visibility. (**B**) Average Ste2 membrane distribution, quantified as in Fig. 6C, in G1 cells with Ste2-sfGFP (blue), Ste2^NPF^-sfGFP (orange), and Ste2^7XR-GPAAD^-sfGFP (green). (**C**) Ste2-sfGFP abundance. Left: representative Western blot. -GFP antibodies label two bands – full-length Ste2-sfGFP and vacuolar sfGFP (note absence of vacuole signal for Ste2^7XR-GPAAD^). Right: quantification of full-length Ste2 abundance (n=3 biological replicates, normalized to the average abundance of wild-type Ste2). (**D**) Halo assay for pheromone sensitivity of cells with wild-type Ste2 (blue), Ste2^NPF^ (orange), and Ste2^7XR-GPAAD^ (green). Top: images of representative halos. Bottom: quantification of halo diameter (n=9, 3 technical replicates at 3 pheromone concentrations, normalized to the average wild-type halo diameter; * t test, p < 0.05). (**E**) Cumulative distribution of the duration of the indecisive phase for MAT**a** cells that were born immediately adjacent to a MATα partner in G1, and either budded or mated by the end of the movie. Cells harboring Ste2 (blue, n=71), Ste2^NPF^ (orange, n=53), or Ste2^7XR-GPAAD^ (green, n=47). (**F**) Left: Cumulative distribution of initial Bem1 cluster orientation relative to the nearest potential mating partner for MAT**a** cells born immediately adjacent to a MATα G1 cell. Cells with wild-type Ste2 (blue, n=117), Ste2^NPF^ (orange, n=78, not significant), or Ste2^7XR-GPAAD^ (green, n=79, not significant). Right: polar histograms of the same data. Scale bar, 3 μm. Strains: DLY20713, DLY20715, DLY21705 (A, B), DLY21203, DLY21206, DLY21704 (C), DLY22321, DLY21301, DLY21295 (D), DLY12943, DLY22058, DLY22397 (E, F).

To quantify the accuracy of initial clustering, we recorded the location of Bem1 clusters among all cells that were born immediately adjacent to a G1 cell of opposite mating type, including those that mated, budded, or failed to do either by the end of the movie. We found that cells despite the dramatic difference in receptor distribution (Fig. 7B), cells with Ste2^NPF^ or Ste2^7XR-GPAAD^ were no less accurate than wild type cells at orienting their initial clusters towards adjacent partners (Fig. 7F). Cells with Ste2^7XR-GPAAD^ were a little more accurate than wild-type cells (Fig 7F), perhaps indicating that abundant and uniformly distributed Ste2 improves gradient sensing, but the difference was not statistically significant for the number of cells analyzed. The finding that even cells with a highly asymmetrical receptor distribution can respond to a pheromone gradient suggests that yeast have a mechanism to correct for variations in receptor density.

### Ratiometric sensing of receptor occupancy

One way to correct for variations in receptor density would be to measure the local ratio of ligand-bound to unbound receptors (i.e. ratiometric sensing). If cells were to respond to the spatial distribution of the *ratio* of active/total receptors, rather than the spatial distribution of active receptors, then differences in the local receptor density would not distort a cell’s ability to determine the orientation of the pheromone gradient. A recent study (Bush, et al., 2016) proposed a mechanism for ratiometric sensing by Ste2, based on the observation that the RGS protein (regulator of G-protein signaling) Sst2 binds to unoccupied Ste2 (Ballon, et al., 2006). Pheromone-bound Ste2 loads GTP on Gα, whereas unbound Ste2-Sst2 promotes GTP hydrolysis by Gα, so the level of activated Gα depends on the ratio between pheromone-bound and unbound Ste2 (Fig. 8A). Although originally proposed as a global mechanism to integrate signaling from all Ste2 (Bush, et al., 2016), in principle this mechanism could also apply locally to extract the gradient of pheromone from the spatial distribution of bound/unbound receptor.

**Figure 8.**
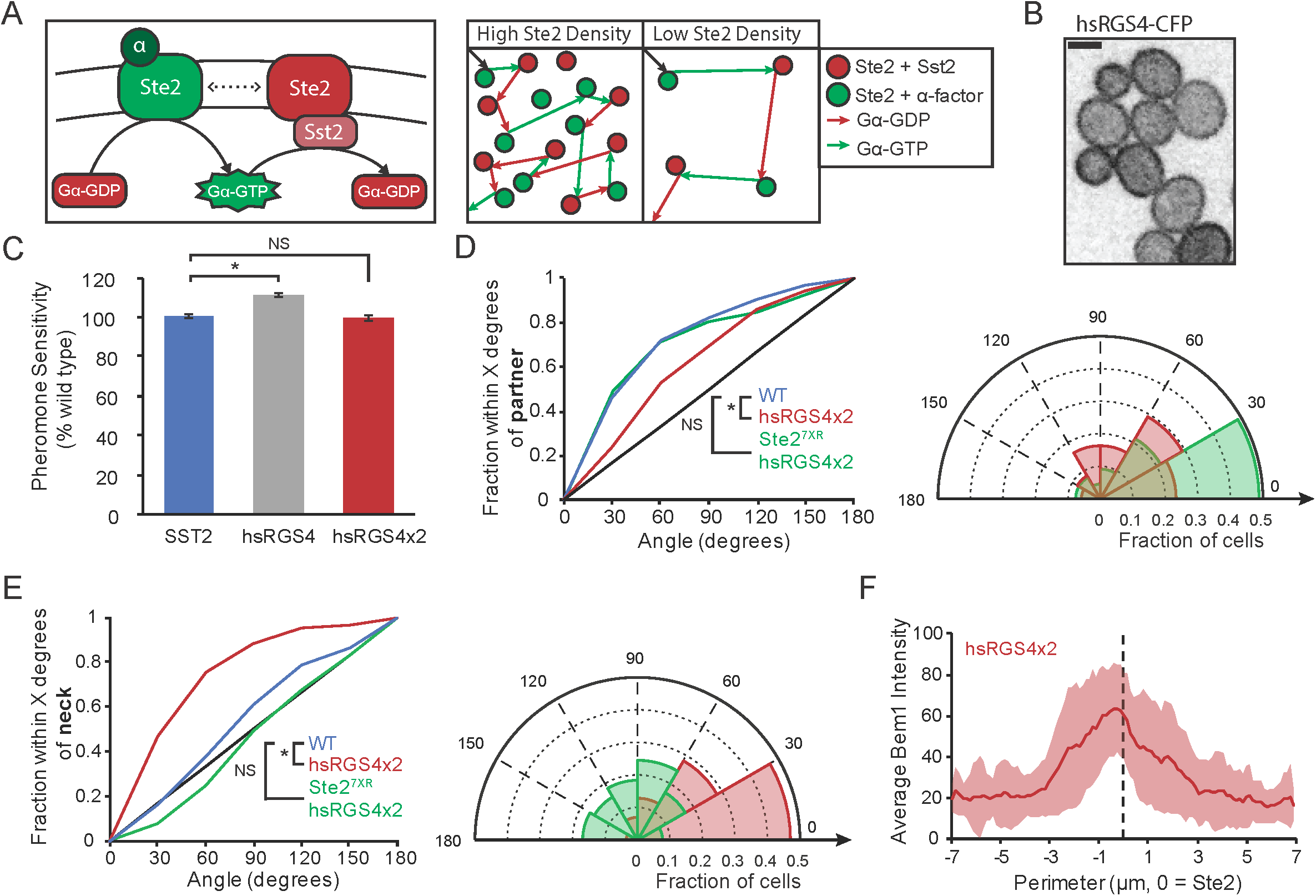
Ratiometric sensing allows cells to orient towards partners despite uneven receptor density. (**A**) Proposed ratiometric pheromone sensing mechanism. Left: Gα is activated by pheromone-bound receptor (Ste2 + α-factor), and inactivated by the RGS protein Sst2. Sst2 associates with inactive Ste2. When Ste2 is activated by α-factor, Sst2 dissociates from Ste2. The instantaneous activation state of Gα is determined by the state of the receptor with which it last interacted. Right: Gα switches between active (green arrows) and inactive (red arrows) states when it interacts with active (green circles) and inactive (red circles) receptors. The fraction of the local Gα that is active reflects the ratio of active to inactive receptors, regardless of receptor density. This means differences in pheromone level at different points on the cell surface can be compared even if there are differences in receptor density. (**B**) hsRGS4 is distributed uniformly on the membrane. Single-plane inverted image of hsRGS4-CFP. (**C**) Pheromone sensitivity measured via halo assay in wild-type cells (blue), and cells in which Sst2 has been replaced by one copy (gray, hsRGS4, * t test, p < 0.05) or two copies (red, hsRGS4×2, not significant) of hsRGS4 (n=9, 3 technical replicates at 3 pheromone concentrations, normalized to the average wild type halo diameter). (**D**) Left: Cumulative distribution of the location of initial Bem1 cluster orientation relative to the nearest potential mating partner in wild type cells (blue, n=117), hsRGS4×2 cells (red, n=85, * two sample KS test, p < 0.05), and hsRGS4×2 cells harboring Ste2^7XR-GPAAD^ (uniform receptor, green, n=65, not significant). Right: polar histogram of the same data for hsRGS4 strains. (E) Left: Cumulative distribution of the location of initial Bem1 cluster formation relative to the site of cytokinesis in wild type cells (blue, n=117), hsRGS4×2 cells (red, n=85, * two sample KS test, p < 0.05), and hsRGS4×2 cells harboring Ste2^7XR-GPAAD^ (uniform receptor, green, n=65, not significant). Right: The same data represented as a polar histogram (WT not plotted).(**F**) Bem initial cluster location is biased by Ste2 distribution in hsRGS4×2 cells. Average Bem1 distribution at the time of initial clustering relative to Ste2 maximum, plotted as in Fig. 6E (n=33). Scale bar, 3 μm. Strains: DLY22318 (B, C), DLY22321, DLY22520 (C-F), DLY12943, DLY22606 (D-F).

Sst2-based ratiometric sensing can be disrupted by replacing Sst2 with a human paralog, hsRGS4, which has similar GAP activity towards Gα but does not associate with Ste2 (Bush, et al., 2016). hsRGS4 is myristoylated and localized uniformly to the plasma membrane (Fig. 8B). We found that two copies of hsRGS4 expressed from the *SST2* promoter were sufficient to restore wildtype pheromone sensitivity (as judged by halo assays) to cells lacking endogenous Sst2 (Fig. 8C). Compared with cells containing Sst2, cells with hsRGS4×2 were significantly worse at orienting at their initial Bem1 clusters towards their partners (Fig. 8D). Instead, the initial clusters in hsRGS4×2 cells were strongly biased towards the previous mother-bud neck (Fig. 8E), a region of high receptor density (Fig. 6). Indeed, a direct comparison showed that unlike wild-type cells, hsRGS4×2 cells displayed a strong tendency to establish initial clusters of Bem1 at sites enriched for Ste2 (Fig. 8F). Thus, gradient sensing depends on the endogenous RGS protein Sst2, which may assist in this process by linking Gα GTP hydrolysis to the location of unbound receptor.

If the inaccurate gradient sensing exhibited by hsRGS4×2 cells is indeed due to disruption of ratiometric sensing, then the orientation defect of hsRGS4×2 should be corrected if the cells were to have uniformly distributed receptors (i.e. ratiometric sensing should be unnecessary if receptor density is uniform). We used the more uniformly distributed Ste2^7XR-GPAAD^ to test this hypothesis, and found that Ste2^7XR-GPAAD^ restored the accuracy of initial Bem1 clustering to wildtype levels in hsRGS4×2 cells (Fig. 8D,E,F). These findings suggest that yeast cells use Sst2-dependent local ratiometric sensing of receptor occupancy to extract accurate information from the pheromone gradient despite having non-uniform receptor density.

## Discussion

### Initial polarity cluster location is surprisingly accurate

The rapid diffusion of peptide pheromones and the small size of the yeast cell led to the expectation that there would be only a small difference in pheromone concentration between the up-and down-gradient sides of the cell. This poses a fundamental difficulty in extracting accurate directional information in the face of molecular noise (Berg and Purcell, 1977). Indeed, a recent study on cells responding to a 0.5 nM/μm pheromone gradient found that initial polarity cluster location was close to random with respect to the gradient (Hegemann, et al., 2015). Moreover, simulations of gradient sensing suggested that even with uniformly distributed receptors and error-free interpretation of the ligand-bound receptor distribution, the signal from such gradients would be obscured by molecular noise and diffusion (Lakhani and Elston, 2017). In principle, time-averaging of the ligand-bound receptor distribution could extract the signal from the noise, but we show that initial clustering of polarity factors occurs 5.1 +/-2.7 min from cell birth, which is too fast to allow for significant time averaging given the slow timescale of yeast pheromone-receptor binding and dissociation (Bajaj, et al., 2004; Raths, et al., 1988; Jenness, et al., 1986). If pheromone levels were high enough, however, this rapid polarization may be able to take advantage of pre-equilibrium sensing and signaling (Ventura, et al., 2014), a proposed mechanism in which directionality is inferred from the rates of pheromone binding rather than steady-state distributions.

Two other factors make it even harder for yeast cells to extract accurate spatial information from pheromone gradients. First, the polarity circuit in yeast contains strong positive feedback sufficient to allow symmetry-breaking polarization in the absence of a directional cue (Chiou, et al., 2017; Johnson, et al., 2011). This allows cells to polarize in random directions when treated with uniform pheromone concentrations (Dyer, et al., 2013; Strickfaden and Pryciak, 2008), and would be expected to enable noise-driven polarization in random directions in cells responding to a shallow pheromone gradient (Chou, et al., 2008). Second, we found that yeast cells do not have uniform receptor distributions. The polarized secretion, slow diffusion, and subsequent endocytosis of pheromone receptors resulted in significant receptor asymmetry, with (on average) three-fold more concentrated receptors on one side of the cell than the other. This creates a receptor gradient that is significantly steeper than the assumed pheromone gradient detected by the cells. As the receptor gradient is randomly oriented with respect to the mating partner, this poses a serious hurdle in accurate gradient detection.

Despite the difficulties enumerated above, we found that the location of initial polarity factor clustering in mating mixtures was highly non-random and surprisingly accurate, with more than 40% of cells clustering within 30° of the correct direction and less than 5% of cells clustering in the opposite segment (a random process would have 17% of cells polarizing in each of these segments). This finding suggests that physiological pheromone gradients may be considerably steeper than previously assumed, and/or that cells possess unappreciated mechanisms to overcome the difficulties in accurate gradient detection discussed above.

### Orientation accuracy is enhanced by ratiometric sensing

One way to avoid being misled by an asymmetric receptor distribution would be to compare the local *ratio* of occupied and unoccupied receptors, rather than simply the density of occupied receptors, across the cell surface. An elegant mechanism to extract information about the fraction of occupied receptors was proposed by Bush, et al. (2016). Because the RGS protein Sst2 binds to unoccupied receptors (Ballon, et al., 2006), those receptors promote GTP hydrolysis by Gα. Conversely, occupied receptors catalyze GTP-loading by Gα. Thus, the net level of GTP-Gα reflects the fraction (and not the number) of occupied receptors on the cell (Bush, et al., 2016). For this mechanism to promote *local* ratiometric sensing requires additionally that a pheromone-bound receptor diffuse slowly relative to its lifetime at the surface (∼10 min) (Jenness and Spatrick, 1986), so that information about where receptors were when they bound to pheromone is not lost. Similarly, GTP-Gα and Gβγ must diffuse slowly relative to the timeframe for Gα GTP hydrolysis and G protein re-association, so that information about where they were when they became activated is not lost.

We found that when RGS function was delocalized by replacing Sst2 (which binds unoccupied receptors) with an equivalent amount of hsRGS4 (which binds the plasma membrane), the accuracy of initial polarity clustering was severely compromised. Instead of polarizing towards potential partners, these cells assembled polarity clusters at regions where receptors were concentrated (often at the site of the last cell division or neck). Thus, abrogating the Sst2-based ratiometric sensing mechanism allowed cells to be misled by the asymmetric receptor distribution. Accurate orientation could be restored to these cells by manipulations that made receptor distribution more uniform. In sum, our findings suggest that local ratiometric sensing compensates for uneven receptor distribution and allows more accurate polarization towards mating partners.

### Why is receptor distribution non-uniform?

Blocking receptor endocytosis allowed receptors to accumulate all over the cell surface in a much more uniform distribution than that seen in wild-type cells. In our mating conditions, this promoted a slightly more accurate orientation of initial polarity factor clustering towards mating partners, and a small improvement in mating efficiency. Why, then, would cells internalize their receptors and create the need for error correction by ratiometric sensing? One possible answer stems from the fact that wild yeast (unlike lab strains) are able to switch mating type. Without receptor endocytosis, cells may be unable to clear pre-existing receptors during mating type switching, generating a situation in which cells arrest in response to their own newly secreted pheromones after a switch. We speculate that receptor endocytosis is necessary to clear the membrane of old receptors when switching mating types, and that receptor asymmetry is the price that cells pay for this advantage.

### Error correction following initial clustering of polarity factors

Although initial polarity clusters were biased to occur near potential mating partners, the process was error-prone and about 60% of cells failed to orient initial polarity within 30° of the correct direction. Nevertheless, these cells did eventually polarize towards partners and mate successfully, indicating the presence of a potent error correction mechanism. We found that after initial clustering, polarity factor clusters relocated erratically during an “indecisive phase” of variable duration (48 +/-24 min). Even cells that had correctly assembled initial polarity clusters close to mating partners exhibited an indecisive phase, although of somewhat shorter duration. During this phase, clusters fluctuated in intensity (concentration of polarity factors in the cluster), extent (broader vs more focused clusters), location, and number (transiently showing no cluster or 2-3 clusters instead of a single cluster). The dynamic polarity clusters were able to polarize actin cables, as we detected frequent accumulation of secretory vesicles at cluster locations. We suggest that this erratic behavior represents a search process in which weak polarity clusters act both as sources of pheromone secretion and locations of pheromone sensing, allowing communication between potential mating partners. At the end of the indecisive phase, cells developed strong and stable polarity sites correctly oriented towards their partners.

The strongest evidence that partners are engaged in communicating with each other during the indecisive phase is that mating pairs ended the indecisive behavior nearly simultaneously (within 5 min of each other). As one partner was born before the other, the durations of their indecisive phases were often quite different, but they transitioned to stable polarization together. Strengthening of the polarity cluster was correlated with an increase in mating MAPK activity, and synthetic induction of MAPK without pheromone led to a similar strengthening of polarity clusters. We speculate that during the indecisive phase, the cells are exposed to a dynamic and constantly changing pheromone landscape. When a mobile polarity cluster is distant from its partner’s cluster, both cells detect relatively low levels of pheromone, leading to intermediate levels of MAPK activity. But if clusters happen to point directly at each other, each cell detects a higher pheromone concentration, leading to an increase in MAPK activity. Increased MAPK then strengthens and stabilizes the polarity cluster, perhaps leading to increased local pheromone secretion and hence increased signaling in a positive feedback loop.

### Exploratory polarization as a mechanism for partner selection

Because the search strategy discussed above depends on polarized pheromone secretion and detection, we call this process “exploratory polarization”. This behavior is strikingly similar to the “speed dating” behavior recently described for mating cells of the distantly related fission yeast *Schizosaccharomyces pombe* (Merlini, et al., 2016; Bendezu and Martin, 2013). In that system, potential mating partners exhibit a prolonged period in which they sequentially assemble and disassemble a weak polarity cluster at multiple locations. Clusters that happen to assemble in the vicinity of a cluster from a mating partner become strengthened and stabilized, presumably due to detection of a higher pheromone level. Thus, distantly related yeasts that mate under very different physiological circumstances (rich nutrients for budding yeast, starvation conditions for fission yeast) appear to have converged on a common and highly effective search strategy.

Exploratory polarization is flexible and responsive to dynamic external conditions. We found that cells abruptly reduced their level of pheromone production when they transitioned from G1 to S phase. Thus, if a potential partner were to enter the cell cycle, a cell would quickly detect reduced pheromone signaling and resume the search for other potential partners. We also noticed that cells with two potential partners nearby could transiently orient clusters towards both partners. However, that situation was unstable and cells only strengthened one polarity cluster and committed to one partner. The basis for restricting polarity to a single site in mating cells is unknown, but may be due to a competition phenomenon as documented for vegetative yeast cells that pick a single bud site (Chiou, et al., 2018; Wu, et al., 2015; Howell, et al., 2012).

An additional elegant feature of exploratory polarization is that it converts a very difficult problem (extracting directional information from shallow and noisy pheromone gradients) into a much easier one (detecting a sharp temporal increase in local pheromone level).

## Conclusions

Yeast cells locate mating partners via a combination of ratiometric spatial sensing and exploratory polarization. Ratiometric sensing of the fraction of occupied receptors compensates for uneven receptor density at the cell surface, allowing cells to decode the pheromone gradient and tentatively identify the locations of potential mating partners. There follows an indecisive period of exploratory polarization in which cells can rapidly scan for partners, wait for nearby potential partners to finish the cell cycle, and identify suitable partners by reciprocal communication through pheromone secretion at the polarity sites. When partners’ polarity clusters align, cells detect higher pheromone levels, leading to increased MAPK activation and stabilization of the polarity site. The partners then enter a committed phase of about 20 minutes with stable polarization and high MAPK activity, after which they fuse.

## Acknowledgements

We thank Serge Pelet (UNIL, Switzerland), Alejandro Colman-Lerner (University of Buenos Aires, Argentina), and Patrick Ferree (Duke University) for providing plasmid reagents. Thanks to Tim Elston, Stefano Di Talia, and Amy Gladfelter as well as members of the Lew lab for stimulating conversations and comments on the manuscript. This work was funded by NIH/NIGMS grants GM103870 and GM122488 to D.J.L.

## Materials and Methods

### Yeast strains and plasmids

Yeast strains used in this study are listed in Table 1. Standard yeast molecular and genetic procedures were used to generate the strains. All strains are in the YEF473 background (*his3-Δ200 leu2-Δ1 lys2-801 trp1-Δ63 ura3-52*) (Bi and Pringle, 1996). The following alleles were previously described: Bem1-GFP (Kozubowski, et al., 2008), Bem1-tdTomato and Spa2-mCherry (Howell, et al., 2012), GFP-Sec4 (Dyer, et al., 2013), STE2^7XR-GPAAD^ (McClure, et al., 2015), Ste7_1-33_-NLS-NLS-mCherry (Durandau, et al., 2015), hsRGS4-CFP (Bush, et al., 2016).

**Table I.**

| **Strain** | **Relevant Genotype** |
| --- | --- |
| DLY7593 | *MATα ura3:BEM1-GFP:URA3* |
| DLY7594 | *MATa ura3:BEM1-GFP:URA3* |
| DLY8156 | *MATα* |
| DLY9070 | *MATα BEM1-GFP:LEU2* |
| DLY12943 | *MATa BEM1-tdTomato:HIS3* |
| DLY13771 | *MATa BEM1-tdTomato:HIS3, GFP-SEC4:URA3* |
| DLY20713 | *MATa SPA2-mCherry:hyg<sup>R</sup>, STE2-sfGFP:URA3* |
| DLY20715 | *MATa SPA2-mCherry:hyg<sup>R</sup>, STE<sup>NPF</sup>-sfGFP:URA3* |
| DLY21070 | *MATα BEM1-tdTomato:HIS3* |
| DLY21203 | *MATa SPA2-mCherry:hyg<sup>R</sup>, STE2-sfGFP:URA3, bar1ΔURA3* |
| DLY21206 | *MATa SPA2-mCherry:kan<sup>R</sup>, STE<sup>NPF</sup>-sfGFP:URA3, bar1ΔURA3* |
| DLY21295 | *MATa STE2<sup>7XR-GPAAD</sup>:URA3* |
| DLY21301 | *MATa STE2<sup>NPF</sup>:URA3* |
| DLY21379 | *MATa BEM1-tdTomato:HIS3, SPA2-GFP:HIS3* |
| DLY21704 | *MATa SPA2-mCherry:kan<sup>R</sup>, STE2<sup>7XR-GPAAD</sup>-sfGFP:LEU2:URA3, bar1ΔURA3* |
| DLY21705 | *MATa SPA2-mCherry:hyg<sup>R</sup>, STE2<sup>7XR-GPAAD</sup>-sfGFP:LEU2:URA3* |
| DLY22058 | *MATa ura3:BEM1-GFP:URA3, STE2<sup>NPF</sup>:URA3* |
| DLY22243 | *MATa BEM1-tdTomato:HIS3, STE2-sfGFP:URA3* |
| DLY22259 | *MATa SPA2-GFP:HIS3, ura3:Ste7<sub>1-33</sub>-NLS-NLS-mCherry:URA3* |
| DLY22318 | *MATa BEM1-GFP:LEU2, SST2:hsRGS4-CFP:kan<sup>R</sup>* |
| DLY22321 | *MATa BEM1-GFP:LEU2* |
| DLY22340 | *MATα BEM1-tdTomato:HIS3* |
| DLY22397 | *MATa BEM1-GFP:LEU2, STE2<sup>7XR-GPAAD</sup>:URA3* |
| DLY22520 | *MATa BEM1-GFP:LEU2, SST2:hsRGS4-CFP:kanR, ura3:P<sub>SST2</sub>-hsRGS4-CFP:URA3* |
| DLY22606 | *MATa BEM1-GFP:LEU2, STE2<sup>7XR-GPAAD</sup>:URA3, SST2:hsRGS4-CFP:kan<sup>R</sup>, ura3:P<sub>SST2</sub>-hsRGS4-CFP:URA3* |
| DLY22883 | *MATα, BEM1-tdTomato:HIS3, leu2:P<sub>MFα1</sub>-sfGFP:LEU2* |
| DLY22764 | *MATa SPA2-GFP:HIS3, ura3:Ste7<sub>1-33</sub>-NLS-NLS-mCherry:URA3, ste5:P<sub>GAL1</sub>-STE5-CTM:P<sub>ADH1</sub>-GAL4BD-hER-VP16:LEU2* |

Spa2-GFP tagged at the endogenous locus was generated by the PCR-based method using pFA6-GFP(S65T)-HIS3MX6 as template (Longtine, et al., 1998).

To express Ste2-sfGFP, sfGFP was amplified by PCR using pFA6a-link-yoSuperfolderGFP-KAN (Addgene plasmid 44901) as template, with primers that added *Not*I sites at the ends. This was used to generate DLB4295, a plasmid with a pRS306 backbone (Sikorski and Hieter, 1989) and a C-terminal piece of the *STE2* ORF (bases 600-1296) fused to sfGFP and followed by 198 bp of the *STE2* 3’-UTR. Digestion at the unique *Cla*I site in *STE2* targets integration of this plasmid to the endogenous *STE2* locus, tagging full-length Ste2 with sfGFP at the C-terminus.

Similar plasmids were used to express Ste2^7XR-GPAAD^-sfGFP (DLB4296) and Ste2^NPF^-sfGFP (DLB4297) at the endogenous *STE2* locus. Ste2^*NPF*^ was generated by first amplifying a fragment of *STE2* using primers that introduced a GGA → AAT mutation (G_392_N substitution) (Tan, et al., 1996) and cloning the fragment back into *STE2*.

Ste7_1-33_-NLS-NLS-mCherry was integrated at *ura3* using pED45 (pRS306-*P*_*RPS2*_*-Ste7*_*1-33*_*-NLS-NLS-mCherry*) as described (Durandau, et al., 2015).

To compromise ratiometric sensing by Sst2, we replaced the endogenous *SST2* with hsRGS4-CFP using A550 (pRS406-K-*hsRGS4-CFP*) as described (Bush, et al., 2016). Because this was insufficient to restore wild-type pheromone sensitivity in our strain background, *P*_*SST2*_*-hsRGS4-CFP* was amplified by PCR and cloned into pRS306 using *Xba*I to generate DLB4414. Digestion with *Stu*I was then used to target integration of a second copy of hsRGS4-CFP at *URA3*.

To make the MFα1 reporter, the MFα1 promoter (506 base pairs upstream of the ATG) was amplified with primers that added *Apa*I and *Hind*III sites, and cloned upstream of a reporter protein with the first 28 residues of Psr1 fused to GFP, followed by the *ADH1* 3’-UTR, in a plasmid with a pRS305 (*LEU2*) backbone (Sikorski and Hieter, 1989). The Psr1_1-28_-GFP reporter was replaced with sfGFP, which was cloned from pFA6a-link-yoSuperfolderGFP-KAN (Addgene plasmid 44901). Digestion at the *PpuM*I in the LEU2 sequence was used to target integration at *leu2*.

To induce MAPK activation without adding pheromone, we generated a plasmid, DLB4239 (pRS305*-*STE5_*5’UTR*_*-* STE5_*3’UTR*_*-* P_ADH1_-GAL4BD-hER-VP16-P_GAL1_-STE5-CTM), that can be used to replace the endogenous *STE5* locus with two genes: (i) a hybrid transcription factor that activates Gal4 target genes in response to estradiol (GAL4BD-hER-VP16) (Takahashi and Pryciak, 2008), and (ii) a *GAL1* promoter driving expression of a membrane-targeted version of Ste5 (P_GAL1_-STE5-CTM) (Pryciak and Huntress, 1998). Addition of estradiol activates the transcription of membrane-targeted Ste5, which leads to activation of the mating MAPKs. The plasmid has a pRS305 (*LEU2*) backbone and contains regions of the *STE5* 5’ and 3’-UTRs upstream of the hybrid transcription factor. DLY4239 was digested with *Pac*I, which cuts between the *STE5* 5’ and 3’-UTR regions to replace endogenous *STE5* with the two genes.

### Live-cell microscopy

Cells were grown to mid-log phase (OD600 ≈ 0.4) overnight at 30°C in complete synthetic medium (CSM, MP Biomedicals, LLC.) with 2% dextrose (Macron). Cultures were diluted to OD_600_ = 0.1. For mating mixtures, the relevant strains were mixed 1:1 immediately before mounting on slabs. Cells were mounted on CSM slabs with 2% dextrose solidified with 2% agarose (Hoefer), which were then sealed with petroleum jelly. For Ste5-CTM MAPK induction (Fig. 4A), slabs also contained 20 nM β-estradiol (Sigma). Cells were imaged in a temperature controlled chamber set to 30°C.

Images were acquired with an Andor Revolution XD spinning disk confocal microscope (Andor Technology, Olympus) with a CSU-X1 5000 rpm confocal scanner unit (Yokogawa), and a UPLSAPO 100x/1.4 oil-immersion objective (Olympus), controlled by MetaMorph software (Molecular Devices). Images were captured by an iXon3 897 EM-CCD camera with 1.2x auxiliary magnification (Andor Technology).

For high resolution images of Ste2-sfGFP, Ste2^NPF^-sfGFP, and Ste2^7XR-GPAAD^-sfGFP (Fig. 6A, C, Fig. 7A, B), z-stacks with 47 planes were acquired at 0.14 μm intervals. The laser power was set to 30% maximal output, EM gain was set to 200, and the exposure for the 488 nm laser was set to 250 ms. For all other microscopy, z-stacks with 15 images were acquired at 0.5 μm z-steps every 2 min, laser power was set to 10% maximal output for the relevant 488 nm, 561 nm, or 445 nm lasers, EM gain was set to 200, and the exposure time was 200 ms.

All fluorescent images were denoised using the Hybrid 3D Median Filter plugin for ImageJ, developed by Christopher Philip Mauer and Vytas Bindokas.

### Analysis of the timing of cell cycle and mating events

Bud emergence was scored using the membrane-targeted Psr1-GFP reporter (Lai, et al., 2018; Kuo, et al., 2014). Cytokinesis was recorded as the first time point when a strong Bem1 signal was visible at the neck. Initial clustering was recorded as the first time point after cytokinesis when a Bem1 cluster was clearly visible and distinguishable from background noise. Polarization was recorded as the time point when the Bem1 patch reached its final stable location and increased in intensity. If the patch appeared at the correct location, but then transiently moved to a new location before returning, polarization was recorded as the time point when the patch returned. Fusion was recorded as the time when cytoplasmic mixing of different color probes became detectable.

### Analysis of polarity factor clustering

To quantify the degree of clustering of the polarity probes Spa2-mCherry, Bem1-tdTomato, and Bem1-GFP, we calculated a “deviation from uniformity” metric from maximum projections of fluorescent z-stack images. Deviation from uniformity, referred to here as clustering (CL), compares the cumulative distribution of pixel intensities in an actual cell, with that in a hypothetical cell with the same range of pixel intensities that are uniformly distributed. That is, CL measures how different the pixel intensity distribution is from a uniform distribution, which reflects the degree to which the signals are clustered.

An elliptical region of interest (ROI) was drawn around each cell at each time point. Raw pixel intensities (p) within each ROI were normalized to a minimum of 0 and maximum of 1:

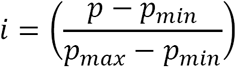

A cumulative distribution (D) of pixel intensities (i) within the cell is then calculated as:

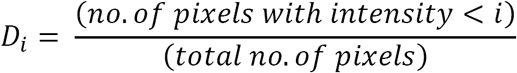

For a cell with uniformly distributed pixel intensities, the cumulative distribution (U) is:

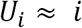

500 uniformly-spaced i-values from 0 to 1 were indexed in ascending order as n = 1, 2, 3,…, 500. The deviation from uniformity metric (CL) was calculated as:

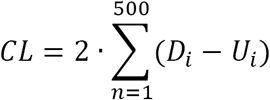

CL approaches a maximum of 1 when a small fraction of pixels exhibit near the maximum intensity, while most pixels are clustered near the minimum intensity – as seen in a highly polarized cell. CL is sensitive to the size of the patch, and the distribution of intensities within the patch – a small patch with sharp edges yields a high CL, while a broad patch with graded edges yields a low CL. As a result, CL is a sensitive indicator of the transition between the indecisive and committed phases.

CL was measured using a MATLAB-based graphical user interface called ROI_TOI_QUANT_V8, developed by Denis Tsygankov.

### Analysis of initial polarity cluster orientation

Initial orientation was measured at the time of initial clustering. For orientation relative to the partner (Fig. 5A,C; Fig. 7F; Fig 8D), we measured the angle between the line from the center of the cell being scored to the centroid of the initial cluster, and a line from the cell center to the closest surface of the nearest G1 cell of the opposite mating type. For orientation relative to the neck (Fig. 5B; Fig. 8E), we measured the angle between the line from the center of the cell being scored to the centroid of the initial cluster, and a line from the cell center to the center of the previous division site. Angles were then grouped into segments of 30° increments.

### Analysis of α-factor synthesis through the cell cycle

The P_MFα1_-sfGFP reporter drives synthesis of sfGFP from the MFα1 promoter. MFα1 is the major α-factor encoding gene. Average fluorescence intensity of the probe was measured from maximum projection images within an elliptical region of interest drawn around each cell. Intensity values were normalized to the value at the end of G1 by dividing by the intensity at the time of bud emergence (for cells with >1 cell cycle, the first bud emergence was used). To express intensity as a function of cell cycle, we set the time of the emergence of the first bud to 0, and the time of the emergence of the second bud to 100.

### Analysis of MAPK activity

MAPK activity was measured using maximum projection fluorescent images of the sensor Ste7-NLS-NLS-mCherry. As demonstrated in (Durandau, et al., 2015), the sensor relocates from the nucleus to the cytoplasm upon phosphorylation by Fus3 or Kss1, and the cytoplasmic to nuclear ratio of the sensor reflects the MAPK activity. We used the coefficient of variation (CV) of pixel intensities measured from maximum projection images to approximate the nuclear to cytoplasmic ratio of the probe. The CV was quite variable from cell to cell, but that variability could be limited by normalization. To approximate MAPK activity (m), an elliptical ROI was drawn around each cell at each time point using ROI_TOI_QUANT_V8. CV was measured for each cell for the 60 minutes prior to fusion, and normalized to a minimum of zero and maximum of 1. Because CV falls as MAPK activity rises was scored as:

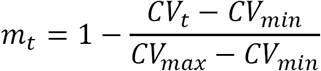

### Analysis of receptor distribution

Membrane distribution of Ste2-sfGFP and Bem1-tdTomato were measured from medial plane fluorescent images. Using FIJI software, fluorescence intensity was averaged across the width of a 3-pixel-wide line tracing the membrane of each cell, drawn with the freehand tool. For comparison of peak location (Fig. 6E; Fig. 8F) the values for individual linescans were normalized by subtracting the background fluorescence, dividing by the maximum point in the linescan, and multiplying by 100 get the %-maximum value. For comparison of receptor uniformity (Fig. 6C; Fig. 7B) the values of individual linescans were normalized by subtracting the background, and bringing each cell to an integral of 1. To generate average distributions, splines were fit to each Ste2 linescan using the smooth.spline function in R, with a 0.75 smoothing factor. The normalized curves for Ste2 or Bem1 from the previous step were then centered on the maximum from the Ste2 spline fit and averaged.

### Halo assays of pheromone sensitivity

Cells were grown to mid-log-phase (OD_600_ ≈ 0.4) at 30°C overnight in YEPD (1% yeast extract, 2% peptone, 2% dextrose). Cultures were diluted to 2.5 x 10^5^ cells/mL, and 5 x 10^4^ cells were spread on YEPD plates in triplicate using sterile glass beads. Plates were allowed to dry for several minutes, and then 2 μL of 1 mM, 500 μM, and 100 μM α-factor was spotted in three separate spots on each plate. Plates were incubated for 48 h at 30°C, and then images were taken using a Bio-Rad Gel Doc XR+ system. Using FIJI software, circles were fit to the zone of arrest surrounding each α-factor spot, and the diameter of the circles was measured in pixels.

### Immunoblotting

Cell cultures were grown in triplicate overnight to mid-log phase in YEPD. 10^7^ cells were collected by centrifugation, and protein was extracted by TCA precipitation as described (Keaton et al., 2008). Electrophoresis and Western blotting were performed as described (Bose et al. 2001). Polyclonal anti-Cdc11 antibodies (Santa Cruz Biotechnology, Inc.) were used at 1:5000 dilution and monoclonal mouse anti-GFP antibodies (Roche) were used at 1:2000 dilution. Fluorophore conjugated secondary antibodies against mouse (IRDye 800CW goat anti-mouse IgG, LI-COR) and rabbit (Alexa Fluor 680 goat anti-rabbit IgG, Invitrogen) antibodies were used at 1:10000 dilution. Blots were visualized and quantified with the ODYSSEY imaging system (LI-COR). All values were normalized to a Cdc11 loading control.

### Statistical analysis

t-Tests were performed in Microsoft Excel via the “t-Test: Two-Sample Assuming Unequal Variances” function (Fig. 7 C, D, Fig. 8C). Two-sample Kolmogorov-Smirnov tests were performed using the Real Statistics Resource Pack software (Release 5.4, developed by Charles Zaiontz) Add-in for Microsoft Excel (Fig. 2D, Fig. 5A-D, Fig. 7F, Fig. 8D, E). p-values over 0.05 were reported as “not significant,” and p-values under 0.05 were reported as “p < 0.05.”

## Figure Legends

**Video 1. Bem1 polarization in a mating mixture.** Cells harboring Bem1-GFP (MAT**α**) and Bem1-tdTomato (MAT**a**) were mixed on an agarose slab and immediately imaged. (Left) False color movie of maximum projection fluorescent images of Bem1-GFP (green) and Bem1-tdTomato (magenta) in a typical mating mixture. (Right) the same movie in inverted grayscale, with labels for budding cells (red dots), G1 phase **α** cells (green dots), G1 phase **a** cells (teal dots), and zygotes (circled in blue). 118 min with 2 min interval between frames. Strains: DLY12943, DLY7593.

**Video 2. Bem1 and Spa2 polarization in mating cells.** MAT**a** cells harboring both Bem1-GFP and Spa2-mCherry were mixed with wildtype MAT**α** cells and immediately imaged. (Top) Maximum projection fluorescent images of Bem1-GFP polarization in 3 example cells from cytokinesis (frame 1) through fusion with a mating partner. (Bottom) Spa2-mCherry polarization in the same three cells. 100 min with 2 min interval between frames. Strains: DLY21379.

**Video 3. MAPK sensor in cycling cells.** The nuclear to cytoplasmic ratio of the MAPK sensor fluctuates through the cell cycle, rising during cytokinesis, and falling during bud growth. Fluorescent images of a field of cells harboring Ste7_1-33_-NLS-NLS-mCherry. 150 min with 2 min interval between frames. Strains: DLY22259.

**Video 4. MAPK sensor in mating cells.** MAPK activity rises (i.e. nuclear to cytoplasmic ratio of the MAPK sensor *falls*) as cells prepare to mate. Fluorescent images of 3 mating type **a** cells harboring Ste7_1-33_-NLS-NLS-mCherry, mating with wildtype mating type **α** cells. 80 min with 2 min interval between frames. Fusion occurs in the final frame for all mating pairs. Strains: DLY22259.

**Figure S1.**
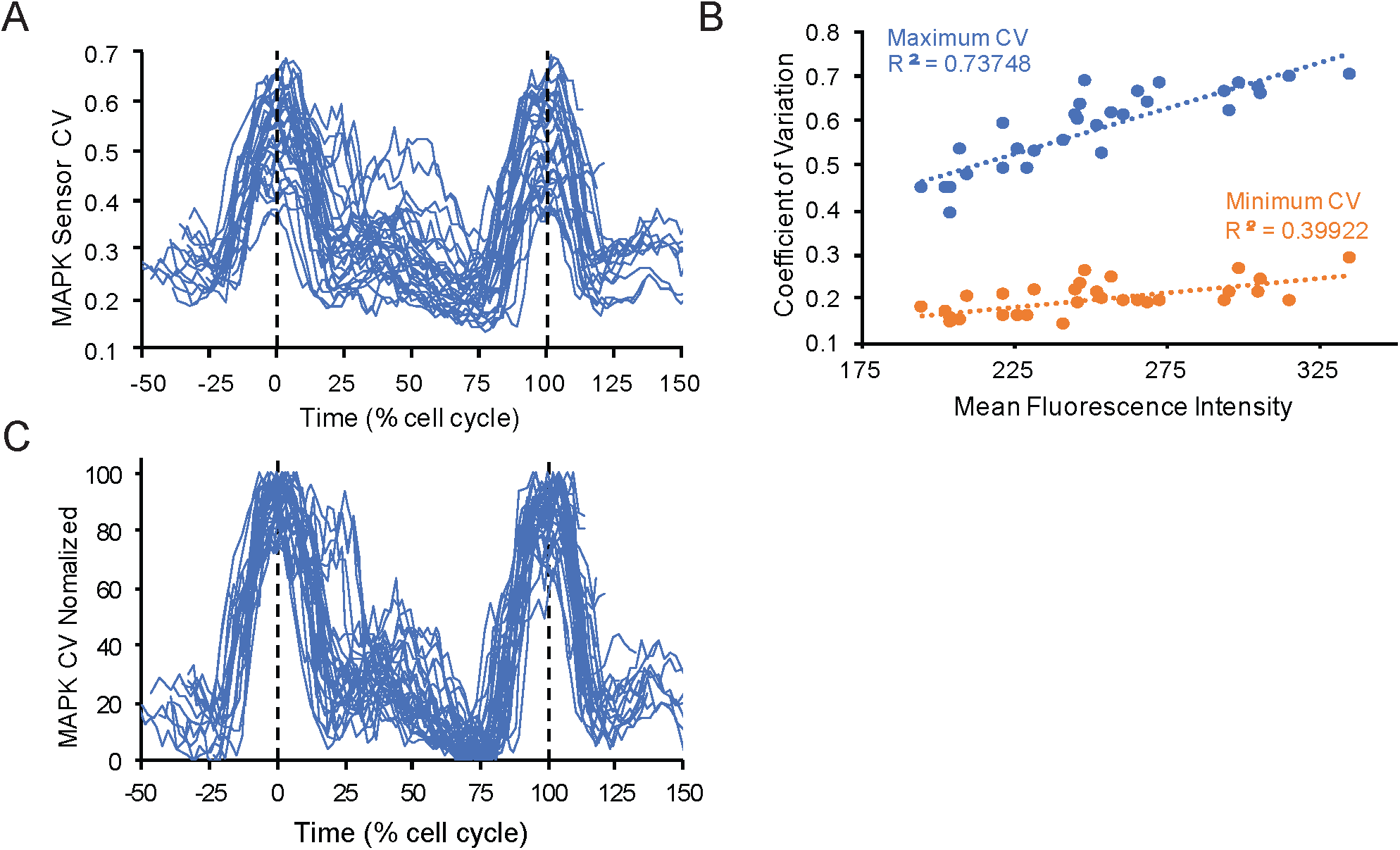
MAPK sensor normalization. Cells harboring Ste7_1-33_-NLS-NLS-mCherry were imaged for 150 min with 2 min resolution. (**A**) Coefficient of variation (CV) of Ste7_1-33_-NLS-NLS-mCherry, measured from maximum projection images in a region of interest encompassing the full cell. Time was normalized to “% cell cycle,” with the first cytokinesis for each cell aligned at 0, and the second cytokinesis aligned at 100. (**B**) Maximum (blue) and minimum (orange) CV vs mean fluorescence intensity for each cell in (A). Mean fluorescence intensity was measured in the same region of interest as CV, and averaged across all time points for each cell. (**C**) Normalized CV, plotted as in (A). CV was normalized to 0 and 1 at the minimum and maximum CV for each cell. Strains: DLY22259 (A-C).

**Figure S2.**
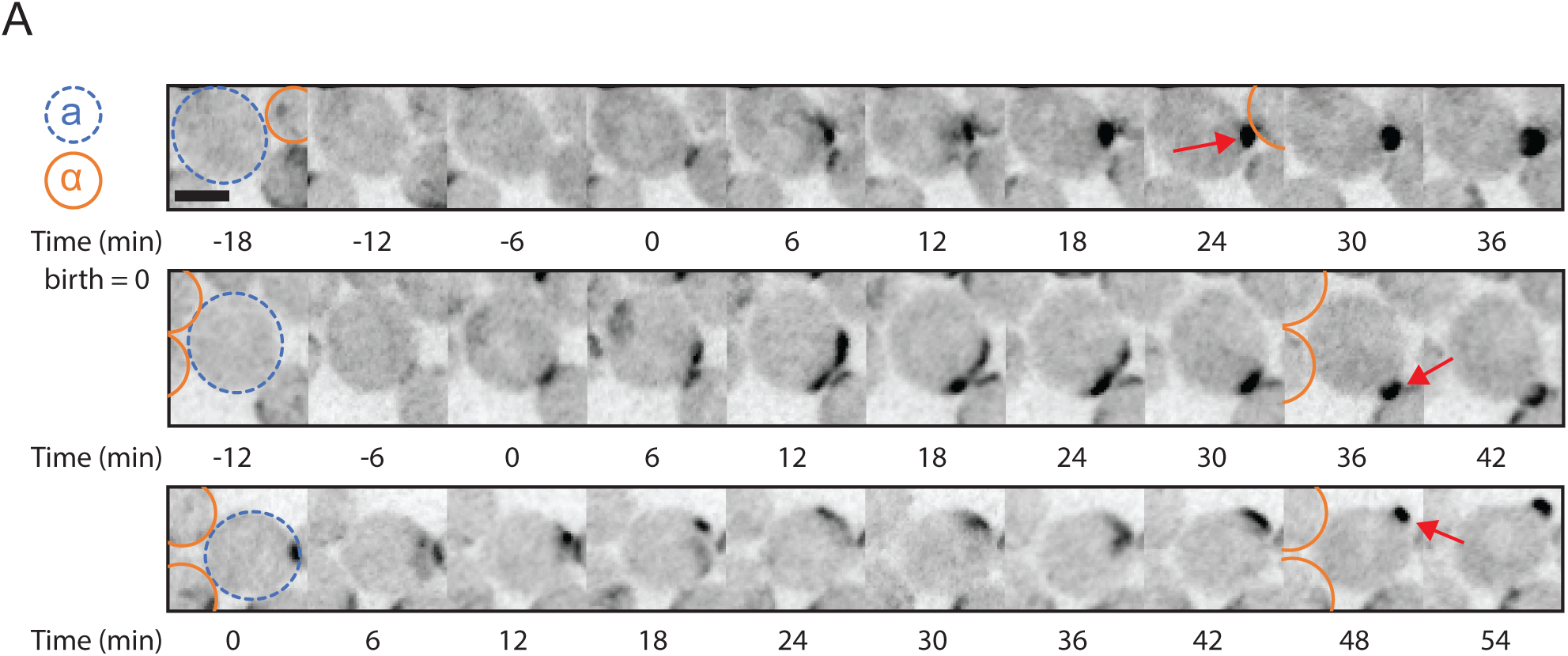
Bem1 polarization in Ste2^NPF^ cells that failed to mate. Cells with faster pheromone receptor endocytosis sometimes bud instead of mating with available partners. MAT**a** cells harboring Bem1-GFP and Ste2^NPF^ were mixed with MATα cells harboring Bem1-tdTomato and imaged immediately. (**A**) Time series of maximum projection images of Bem1-GFP polarization in three MAT**a** cells (blue circles) harboring Ste2^NPF^ that failed to mate with adjacent G1 MATα cells (orange circles). Red arrows indicate bud emergence. 60 min, 6 min interval, birth = 0 min, scale bar, 3 μm. Strains: DLY22058.

